# Temporal Single Cell Analysis of Leukemia Microenvironment Identifies Taurine-Taurine Transporter Axis as a Key Regulator of Myeloid Leukemia

**DOI:** 10.1101/2024.05.11.593633

**Authors:** Benjamin J. Rodems, Sonali Sharma, Cameron D. Baker, Christina M. Kaszuba, Takashi Ito, Jane L. Liesveld, Laura M. Calvi, Michael W. Becker, Craig T. Jordan, John M. Ashton, Jeevisha Bajaj

## Abstract

Signals from the microenvironment are known to be critical for development, sustaining adult stem cells, and for oncogenic progression. While candidate niche-driven signals that can promote cancer progression have been identified^1–6^, concerted efforts to comprehensively map microenvironmental ligands for cancer stem cell specific surface receptors have been lacking. Here, we use temporal single cell RNA-sequencing to identify molecular cues from the bone marrow stromal niche that engage leukemia stem cells (LSC) during oncogenic progression. We integrate these data with our RNA-seq analysis of human LSCs from distinct aggressive myeloid cancer subtypes and our CRISPR based *in vivo* LSC dependency map^7^ to develop a temporal receptor-ligand interactome essential for disease progression. These analyses identify the taurine transporter (TauT)-taurine axis as a critical dependency of myeloid malignancies. We show that taurine production is restricted to the osteolineage population during cancer initiation and expansion. Inhibiting taurine synthesis in osteolineage cells impairs LSC growth and survival. Our experiments with the TauT genetic loss of function murine model indicate that its loss significantly impairs the progression of aggressive myeloid leukemias *in vivo* by downregulating glycolysis. Further, TauT inhibition using a small molecule strongly impairs the growth and survival of patient derived myeloid leukemia cells. Finally, we show that TauT inhibition can synergize with the clinically approved oxidative phosphorylation inhibitor venetoclax^8, 9^ to block the growth of primary human leukemia cells. Given that aggressive myeloid leukemias continue to be refractory to current therapies and have poor prognosis, our work indicates targeting the taurine transporter may be of therapeutic significance. Collectively, our data establishes a temporal landscape of stromal signals during cancer progression and identifies taurine-taurine transporter signaling as an important new regulator of myeloid malignancies.

## INTRODUCTION

Signals from the tumor microenvironment (TME) can regulate the initiation, progression, and immune evasion of tumors^10–16^. For instance, Wnts from immune and stromal cells^2, 3, 17^, integrin ligands on endothelial cells^1^ and, HGF and CXCL12 from myofibroblasts^4–6^, influence tumor growth and therapy resistance. While recent single cell based studies have identified the cellular TME components, especially in the context of solid tumors^18, 19^, concerted efforts to link ligands from the changing TME landscape with cognate cancer cell surface receptors have been lacking. Given that cell surface signals are particularly amenable to therapeutic targeting, functional characterization of these interactions as the disease progresses is of significant clinical interest.

Aggressive therapy resistant myeloid cancers, such as blast crisis phase chronic myeloid leukemia (bcCML) and acute myeloid leukemia (AML), initiate and expand in a complex bone marrow microenvironment. While recent work has described the cellular composition of the normal bone marrow niche^20–22^, dynamic alterations within these populations during disease initiation and establishment remain undefined. We use single cell RNA-sequencing to establish temporal changes in the bone marrow microenvironmental populations and in niche-driven signals during disease progression. To define TME ligands essential for leukemogenesis, we focused on cognate cancer cell surface receptors enriched in both human AML and bcCML stem cells as compared to healthy controls in our new patient dataset, and established as key *in vivo* leukemia stem cell dependencies^7^. This approach identified signals known to be critical for cancer growth, such as Kit/KitL^23^ and Cd47/Thrombospondin1^24^, as well as multiple new signaling axes.

Of these, TauT encoded by the *SLC6A6* gene, was strongly associated with poor prognosis in human leukemias and emerged as a key new regulator of aggressive myeloid cancers. Although taurine supplements are thought to be neuroprotective and often used to mitigate the side-effects of chemotherapy^25–27^, a cancer promoting role of taurine has not been considered. Here, we determine if blocking taurine production from the TME impairs leukemia stem cell function. We also use genetic and small molecule inhibitor-based approaches to establish whether TauT expression in cancer cells controls the initiation and progression of aggressive myeloid leukemias.

## RESULTS

### Temporal changes in myeloid leukemia bone marrow microenvironmental populations

To define temporal changes in the non-immune bone marrow microenvironmental populations during myeloid leukemia progression, we focused on the BCR-ABL/NUP98-HOXA9-driven model of bcCML, which can engraft and grow in an unirradiated healthy murine bone marrow microenvironment. We first established a time course of disease progression based on median survival and identified four distinct time intervals that correlated with different stages of disease progression: naïve, initiation, expansion, and end point (Fig. S1a). Bone marrow stromal populations (leukemia^-^Cd45^-^Ter119^-^) were isolated from leukemic cohorts at each stage and analyzed by single cell RNA-Sequencing (Fig 1a). Clustering the stromal cells based on gene expression resulted in twenty-one lineage clusters with transitional subsets covering endothelial cells (three arteriolar and two sinusoidal subsets expressing *Pecam1* and *Cdh5*), chondrocytes (five subsets expressing *Col2a1* and *Acan*), fibroblasts (four subsets expressing *S100a4*), pericytes (one subset expressing *Acta2*), mesenchymal stromal cells (MSC; expressing *Lepr*), and osteo-associated populations (five subsets expressing *Bglap* and *Col1a1*; Fig. 1b-c).

**Figure 1:**
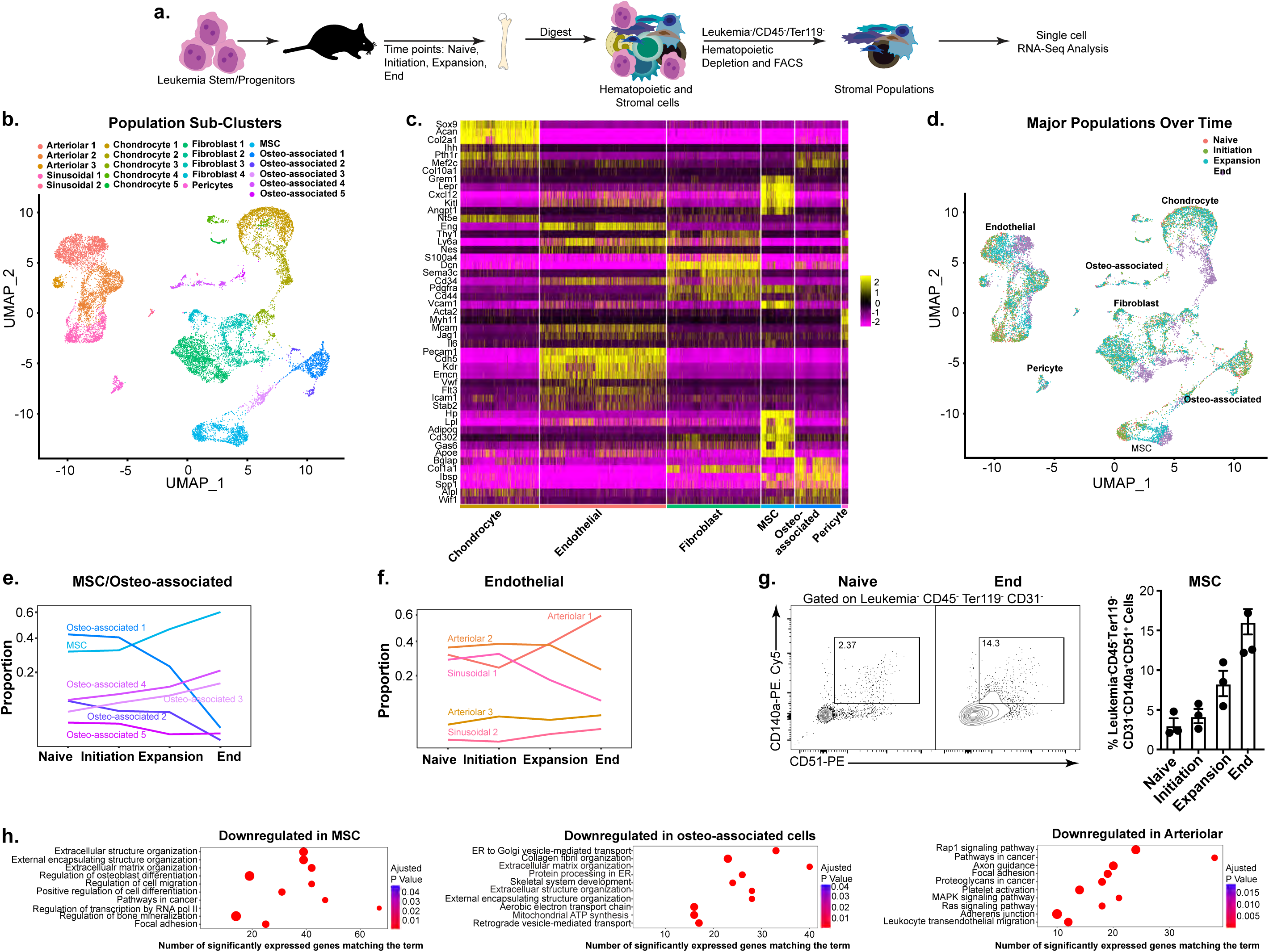
Temporal scRNA-Seq Analysis of Myeloid Leukemia Bone Marrow Microenvironment. (a) Schematic shows the strategy used to determine the impact of bcCML on the bone marrow microenvironment. (b) UMAP of 15695 nonhematopoietic cells (mixed bone and bone marrow, n=3-5 mice/time point) colored by population cluster shows the presence of 21 distinct bone marrow stromal cell clusters. (c) Heatmap of significantly expressed marker genes for nonhematopoietic bone marrow populations. (d) UMAP plot of population clusters over time (naïve=0 days, initiation=2-4 days, expansion=7-9 days and, end=11-14 days post-transplant; Colors represent different stages of disease). (e-f) Line graphs show proportion of mesenchymal stromal cells/osteo-associated (e) and endothelial cells (f) over time. (g) Representative FACS plots and quantification of MSC frequency (Leukemia^-^CD45^-^Ter119^-^CD31^-^CD140a^+^CD51^+^) over time (n=3 mice per timepoint). (h) Enrichr plots show top fifteen downregulated pathways by population cluster in MSCs, osteo-associated, arteriolar endothelial, and sinusoidal endothelial cell populations.

Our temporal analysis identified significant remodeling of the bone marrow microenvironment with disease progression (Figs. 1d and S1b). While MSCs, arteriolar endothelial cells and the immature osteo-associated subsets 3 and 4 expanded, the more mature osteo-associated clusters 1 and 2, and sinusoidal endothelial subset 1 were lost as the disease progressed (Figs. 1e-f). Our flow cytometry-based analyses of an independent cohort of leukemic mice showed a similar increase in MSCs and arteriolar endothelial cells (Figs. 1g and S1c) with a concomitant decline in the sinusoidal endothelial populations (Fig. S1c), validating our scRNA-seq analysis. Clustering analyses of genes expressed in the major lineages identified multiple gene sets with similar changes in expression pattern during disease progression (Fig. S1d), suggesting that these may represent altered stromal cell fate. Consistent with this, pathway analysis showed that osteoblast differentiation and bone mineralization were downregulated in MSCs, and skeletal system development was downregulated in osteo-associated cells with disease progression (Fig. 1h). These data confirmed the phenotypic arrest in osteogenic differentiation and expansion of MSCs in the leukemia niche (Fig. 1e, g). Our analysis also identified a loss in focal adhesion, adherens junctions, and cell migration in endothelial populations (Figs. 1h and S1e), indicating that endothelial cell integrity and function may be impaired with disease progression. Collectively, our temporal analysis of the TME identifies dynamic changes within stromal sub-populations during leukemia progression.

### Mapping leukemia stem cell receptor interactions with microenvironmental ligands

Our experiments above establish distinct temporal changes in the major bone marrow stromal cell lineages in response to leukemia progression. To define the functional relevance of these microenvironmental changes on leukemogenesis, we focused on cell surface signals expressed on leukemia stem cells that act as receptors for niche-driven signals. To identify cancer cell surface receptors associated with disease progression, we carried out RNA-sequencing based gene expression in human AML and bcCML CD34^+^ leukemia stem-enriched cells (LSCs), and healthy donor bone marrow CD34^+^ hematopoietic stem and progenitor cells (HSPCs). Our data shows distinct clustering of normal bone marrow, with some overlap between bcCML and AML samples (Fig. 2a). In these data, there was a significant (P_adj_<0.05) upregulation of 1,569 genes in bcCML, 2,842 genes in AML, and 2,331 genes in both bcCML and AML compared to normal HSPCs. To identify cell surface receptors that are functionally relevant for disease progression, we focused on cell surface genes^28^ that drop-out by two-fold or more in our genome-wide *in vivo* leukemia CRISPR screen^7^. This analysis identified 13 cell surface signals common to both bcCML and AML, 18 unique to AML, and 7 unique to bcCML (Fig. S2a). Of these 38 genes, 16 were misannotated as cell surface in the reference dataset^28^ and removed from further analysis (Fig. 2b, see methods for excluded gene list).

**Figure 2:**
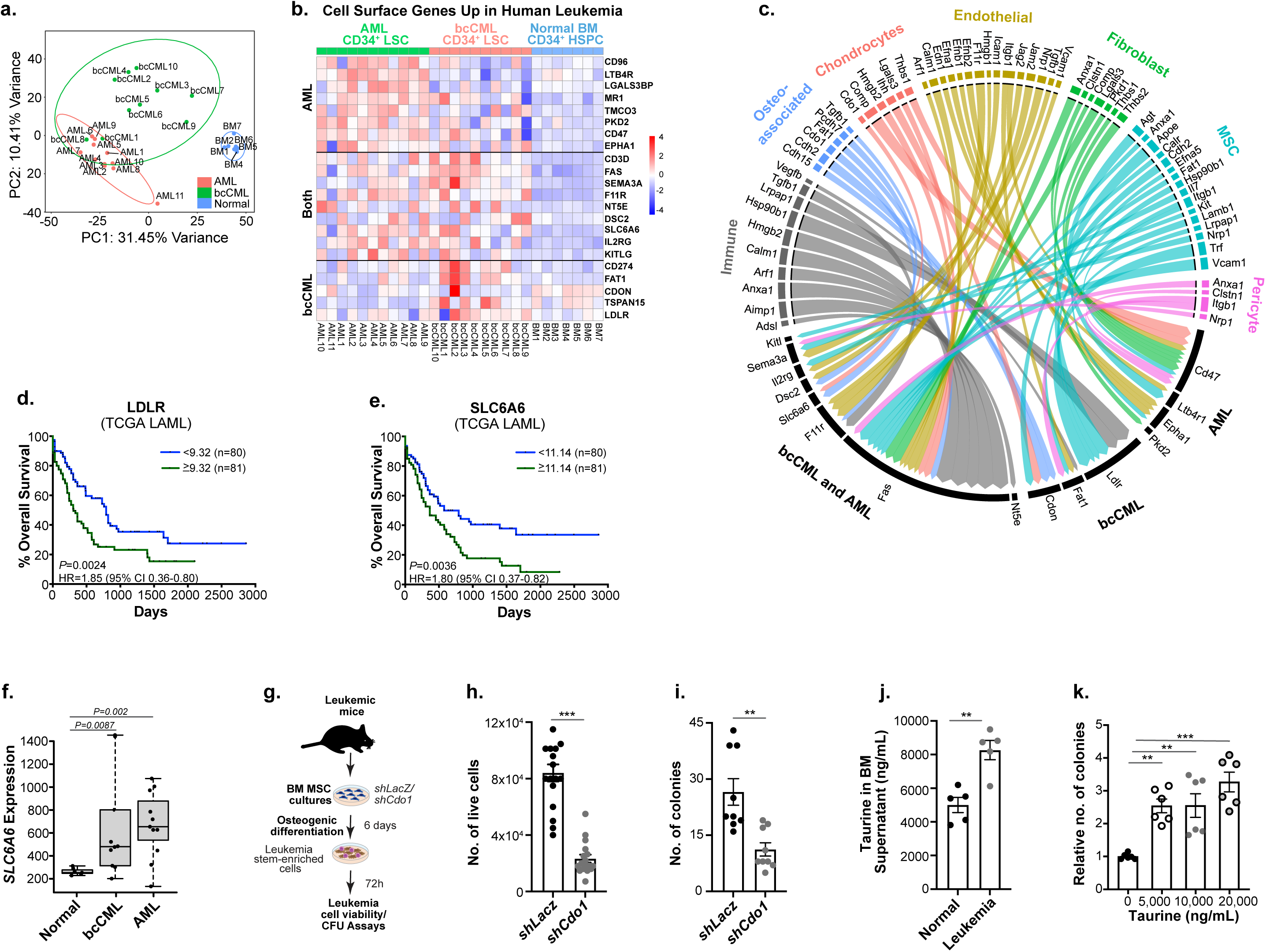
Bone Marrow Microenvironmental Ligands for LSC Specific Cell Surface Receptors. (a) PCA-plot shows the distribution of seven CD34^+^ healthy donor bone marrow hematopoietic stem progenitor cells, ten bcCML CD34^+^ LSCs, and eleven AML CD34^+^ LSCs from human samples (n=7 BM, n=10 bcCML, n=11 AML). (b) Heatmap shows r-log normalized RNA expression of cell surface receptors upregulated in AML, bcCML or both. (c) Circos plot of cognate leukemia cell surface receptor-stromal cell ligand pairs. (d-e) Overall survival of human leukemia patients with (d) high (<9.32, n=80) or low (≥ 9.32, n=81) LDLR expression and (e) high (<11.14, n=80) or low (≥11.14, n=81) SLC6A6 expression (TCGA-LAML; Xena Browser). (f) Normalized *SLC6A6* expression in normal human BM (CD34^+^), bcCML, and AML patient samples (n=7 BM, n=10 bcCML, n=11 AML). (g) Schematic shows the experimental strategy used to determine the impact of inhibiting *Cdo1* in leukemic BM MSCs on co-cultured LSCs. MSCs isolated and cultured from leukemic mice were transduced with either control (sh*LacZ*) or Cdo1 (sh*Cdo1*) lentiviral shRNAs. Six days post induction of osteogenic differentiation, Lin-bcCML LSCs were added to these cultures and LSC survival and colony forming ability was assessed after 72h of co-culture with osteolineage cells. (h-i) Number of live LSCs (h), and their colony forming ability (i) post coculture with leukemic MSCs transduced with *shCdo1* or *shLacZ* (data combined from 3 independent experiments). (j) Taurine quantity per femur in control and leukemic mice, twelve days post-transplant (n=5 per cohort; data combined from two independent experiments). (k) Relative colony forming ability of primary human bcCML cells in the presence of water (control) or indicated amounts of taurine, (n=3 independent culture wells per cohort; data combined from two independent experiments; one-way ANOVA). Error bars represent ±SEM; Statistics from unpaired t-tests. *p*<*0.05, **p*<*0.01, ***p*<*0.001

We next determined the ligands for these 22 cell surface genes upregulated in human AML/bcCML LSCs that are critical for LSC growth *in vivo*, using NicheNet^29^ and literature research (Fig. S2a). For these 22 receptors, we identified ligands significantly upregulated in our leukemia TME scRNA-Seq and in a recent human AML immune microenvironment dataset^30^ (Fig. S2b) at each time point. This analysis led to the exclusion of receptors with no known ligands (*MR1*, *TMCO3* and *TSPAN15*) as well as those whose ligands were not significantly enriched in any TME population (*LGALS3BP*, *CD96*, *CD274*, and *CD3D*). We thus generated a unique map of TME ligands for 15 LSC cell surface receptors essential for disease progression, that included interactions known to be critical for cancer growth such as Kit/KitL^23^ and Cd47/Thrombospondin1^24^ (Fig. 2c).

In addition to identifying key ligand-receptor interactions, we sought to determine whether disease progression can alter the expression of LSC-specific TME ligands. Our analysis identified four distinct patterns of ligand expression. While the expression of some genes remained steady at all time points, e.g., *Jam2* in arteriolar endothelial cells and *Ihh* in the chondrocytes (Fig. S2c), the expression of genes such as *Vcam1*, *Pcdh7* and *Il7* was lost in the TME, especially during disease end point (Fig. S2d). Expression of genes like *Anxa1* first increased during disease progression but then declined towards the end (Fig. S2e). While a fourth category of genes were expressed at all time points, the microenvironmental populations expressing them changed over time. These included *Pvr*, *Cdo1*, *Apoe* and *Agt* (Fig. S2f). Our temporal analysis thus identifies signals that are likely necessary not only at different stages of disease progression, but also those that are critical during the entire course of the disease.

### Functional Characterization of TME Ligands essential for Leukemia Progression

While all the TME-LSC signaling axes identified in our interactome are likely to be essential for leukemia progression, we focused on genes associated with unfavorable outcomes in human AML patients. Of the 22 genes upregulated in human LSCs (Fig. 2b-c), only *LDLR* (Fig. 2d, Fig. S3a) and *SLC6A6* (Fig. 2e-f) were significantly associated with poor prognosis in AML (TCGA-LAML). We thus tested the functional requirement of both LDLR and SLC6A6 ligands from the microenvironment on bcCML stem cell growth and survival in co-culture assays.

One of the primary Low-Density Lipoprotein Receptor (LDLR) ligands, Apolipoprotein E (ApoE)^31, 32^, was highly expressed in MSCs at all time points (Fig. S2f). To test the role of ApoE from MSCs on leukemia stem cell function, we co-cultured bcCML lineage^-^ leukemia stem-enriched cells (LSCs) with MSCs transduced with shRNAs targeting *ApoE* or *LacZ* (control; Fig. S3b). Co-culture with MSCs lacking *ApoE* expression (Fig. S3c) led to a significant reduction in both the viability and colony forming ability of bcCML LSCs (∼2.1-fold lower than controls; Fig. S3d-e). These data validate our approach to identify TME-ligands for LSC surface receptors and indicate a critical role of MSC-driven ApoE expression in sustaining LSC survival and self-renewal.

*SLC6A6* encodes the taurine transporter (TauT) and is required for the cellular uptake of the non-essential amino acid taurine. Although TauT can also transport β-alanine, it has 12.4-fold stronger affinity for taurine (taurine Km=4.5μM vs. β-alanine Km=56μM)^33^. While our analyses of the leukemia TME identified significant expression of enzymes necessary for taurine biosynthesis (Cdo1 and Csad)^34^, those required for β-alanine synthesis (Gadl1 and Cndp1)^35, 36^ could not be detected. These data indicate that taurine is likely the key TauT ligand synthesized by the bone marrow niche. Given that *Cdo1* expression was restricted to the osteo-associated cells during cancer initiation and expansion (Fig. S2f; Fig. S3g), the osteolineage populations are likely the key source of taurine in the TME. To test this, we determined *Cdo1*/*Csad* expression and taurine production during MSC osteogenic differentiation (Fig. S3h). Our data showed a striking increase in *Cdo1* and *Csad* levels at day 6 (Fig. S3i), accompanied with a 2.3-fold increase in taurine levels in culture media, confirming that osteolineage cells can indeed produce taurine (Fig. S3j). To determine a functional role of taurine from the osteolineage cells we tested the impact of inhibiting taurine production on co-cultured LSCs (Fig. 2g). Our experiments showed that LSCs co-cultured with osteolineage cells lacking *Cdo1* expression (Fig. S3k) were significantly less viable (3.6-fold reduction compared to controls; Fig. 2h) and formed fewer colonies (2.4-fold lower than controls; Fig. 2i). These results further validate our functional strategy to identify key niche driven ligands essential for LSC function. Consistent with a functional role of taurine from the TME in leukemia progression, we saw 1.7-fold higher taurine levels in leukemic bone marrow as compared to controls (Fig. 2j). Moreover, exogenous taurine supplements increased LSC colony forming ability by 2.5 to 3.3-fold (Fig. 2k).

Collectively, these data indicate a key role of taurine and ApoE from the bone marrow niche in sustaining LSC growth and survival. Unlike *LDLR*, *SLC6A6* expression was significantly enriched in both bcCML and AML LSCs as compared to controls (Fig. 2f, S3a). We thus focused our studies on *SLC6A6* since it is likely that this gene may be broadly required for the growth of aggressive myeloid leukemias.

### TauT in normal hematopoiesis

To test the role of TauT in normal hematopoiesis using definitive genetic approaches, we used the global TauT knockout mice (Fig. 3a)^37^. While TauT knockout mice are born in Mendelian ratios, they can develop ageing related defects in bone mass and retinal degeneration^37–39^. Consistent with a requirement for TauT expression for taurine uptake, taurine levels in the TauT^-/-^ bone marrow cells were reduced by 5.7-fold compared to wild type controls (Fig. 3b). An analysis of the TauT^-/-^ mice indicated that its loss did not impact bone marrow cellularity (Fig. 3c), or the total numbers of HSCs, MPPs (Figs. 3d-e) and the committed GMP, CMP, MEP progenitors (Fig. 3f-g). However, there was minor decrease in the frequency of MPPs (Fig. S4a) and CMPs (Fig. S4b) and an increase in MEPs (Fig. S4b). Although we noted a trend towards increased Ter119^+^ erythroid cells consistent with the increase in MEPs (Figs. 3g-h and S4b, d), there was no significant difference in the frequency or number of differentiated populations (Figs. 3h and S4c-d). While an analysis of the peripheral blood similarly showed no significant changes in most complete blood cell counts (Figs. S4 e-j), we identified a small reduction in platelets with TauT loss (Fig. S4h).

**Figure 3:**
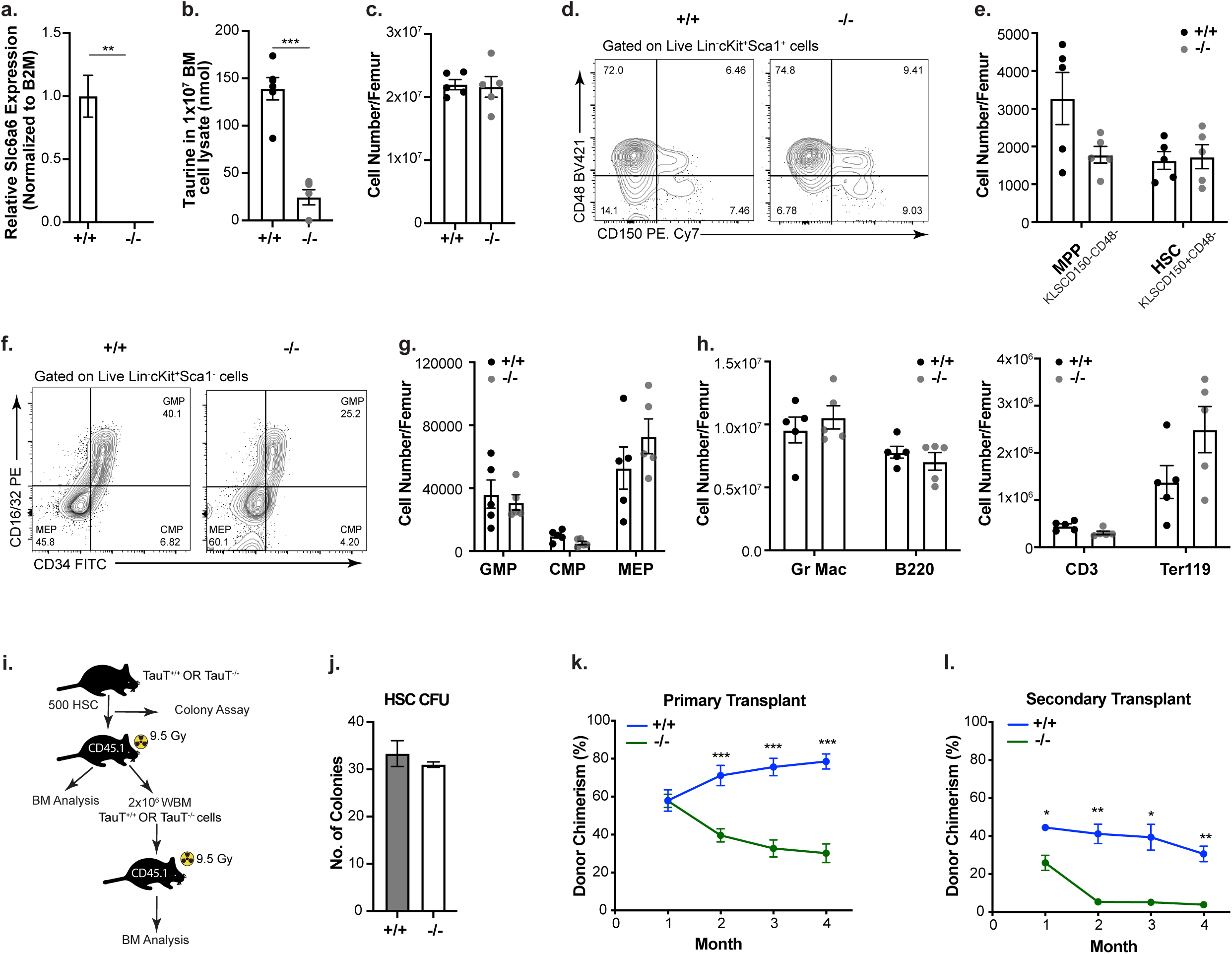
The Role of TauT in Normal Hematopoiesis. (a) Relative *Slc6a6* mRNA expression in whole bone marrow of +/+ (TauT^+/+^) and -/- (TauT^-/-^) mice (n=3 technical replicates per cohort). (b) Taurine quantity in whole bone marrow cell lysate of +/+ and -/- mice (n=6 pelvic bones from 3 mice per cohort; data combined from two independent experiments). (c) Average number of bone marrow cells in +/+ and -/- mice. (d) Representative FACS plots show frequency of HSC and MPP cell numbers in +/+ and -/- mice (gated on Lin^-^cKit^+^Sca^+^ or KLS cells, where Lineage^+^ is marked by CD3e, CD4, CD8, Gr1, CD11b/Mac-1, Ter119, B220 and CD19). (e) Average number of HSCs (KLSCD150^+^CD48^-^) and multipotent progenitors (MPPs, KLSCD150^-^CD48^-^) in +/+ and -/- mice. (f, g) Representative FACS plots (f) and quantification of average number of committed progenitors (granulocyte–macrophage progenitors or GMP, Lin^−^IL7Ra^−^Kit^+^Sca1^−^CD34^+^CD16/3^+^; common myeloid progenitors or CMP, Lin^−^IL7Ra^−^Kit^+^Sca1^−^CD34^+^CD16/32^−^; megakaryocyte–erythroid progenitors or MEP, Lin^−^IL7Ra^−^Kit^+^Sca1^−^CD34^−^CD16/32^−^) in +/+ and -/- mice. (h) Total number of differentiated hematopoietic cells in bone marrow of +/+ and -/- mice (For c-h, n=5 from age and sex matched littermates; data combined from four independent experiments). (i) Schematic shows the experimental strategy to determine the impact of TauT loss on the *in vivo* repopulating capacity and self-renewal ability of HSCs. BM, bone marrow; WBM, whole bone marrow. (j) Colony-forming ability of +/+ and -/- HSCs (n=3 independent culture wells per cohort). (k, l) Graphs show average donor chimerism in the peripheral blood of primary and secondary transplant mice (Primary: n=9-10 per cohort for primary transplants; data combined from two independent experiments. Secondary: n=6 per cohort). Error bars represent ±SEM; statistics from unpaired t-tests or as indicated. *p*<*0.05, **p*<*0.01, ***p*<*0.001.

Functionally, TauT deletion did not impact HSC colony formation (Figs. 3i-j). While TauT loss did not impair initial HSC engraftment (1 month post-transplant), donor chimerism dropped over time (Fig. 3k). Although the overall bone marrow chimerism 4 months post-transplant was 2.4-fold lower in TauT^-/-^ HSC recipients as compared to TauT^+/+^ (Fig. S4k), these TauT^-/-^ HSCs were able to contribute significantly to all hematopoietic lineages (Figs. S4l-n). Serial transplantation of TauT^+/+^ and TauT^-/-^ bone marrow cells showed a similar loss in TauT^-/-^ engraftment over time (Figs. 3l and S4o-s). These results suggest that while TauT loss may not impair steady-state hematopoiesis, it can impact long term self-renewal and maintenance. These data are similar to ours and others’ prior findings with genetic loss of LSC enriched genes on normal HSC function, including CD98^1^, Staufen 2^7^, Musashi 2^40^, Brd4^41, 42^, and Bcl2^43^. However, therapeutic inhibition of CD98 showed minimal toxicity in AML Phase I trials^44^ and Bcl2 inhibitors are approved for AML therapy^8^. It is thus likely that small molecule inhibitor based approaches may identify a therapeutic window for TauT targeting in human cells.

### TauT loss impairs myeloid leukemia initiation and depletes leukemia stem cells

Consistent with a critical requirement for taurine from the bone marrow niche in promoting leukemia growth (Fig. 2h), the loss of its transporter TauT significantly increased the median survival of mice transplanted with primary bcCML (3.9-fold higher likelihood of survival, Figs. 4a-b). TauT loss also led to a functional depletion in the bcCML stem cell population as indicated by a 2.3-fold reduction in the *in vitro* colony forming ability (Fig. 4c), as well as a striking increase in survival of mice transplanted with TauT^-/-^ LSCs (40%) relative to control (0%) in secondary transplant assays (21.2-fold higher likelihood of survival, Fig. 4d). The 8.5-fold reduction in the *in vitro* colony-forming ability of bcCML LSCs from secondary transplants (Fig. 4e) suggests that these leukemias remain dependent on TauT for their continued propagation. To determine if TauT is broadly required for *de novo* AML growth, we tested the impact of its loss on the initiation of leukemia driven by MLL-AF9/ NRAS^G12V^ as well as a model driven by AML-ETO9a/NRAS^G12V^. Our experiments show that TauT loss markedly delays the initiation of MLL-driven AML relative to control (6.1-fold higher likelihood of survival, Figs. 4f-g). TauT^-/-^ cKit^+^ AML stem cells from established disease formed 2.9-fold less colonies compared to controls (Fig. 4h), indicating that TauT loss depleted functional leukemic stem cell populations. Consistent with a key role of TauT expression in myeloid leukemia initiation, TauT loss in disease driven by AML-ETO9a resulted in a striking increase in survival (70%) relative to controls (0%) (36.3-fold higher likelihood of survival, Fig. 4i-j).

**Figure 4:**
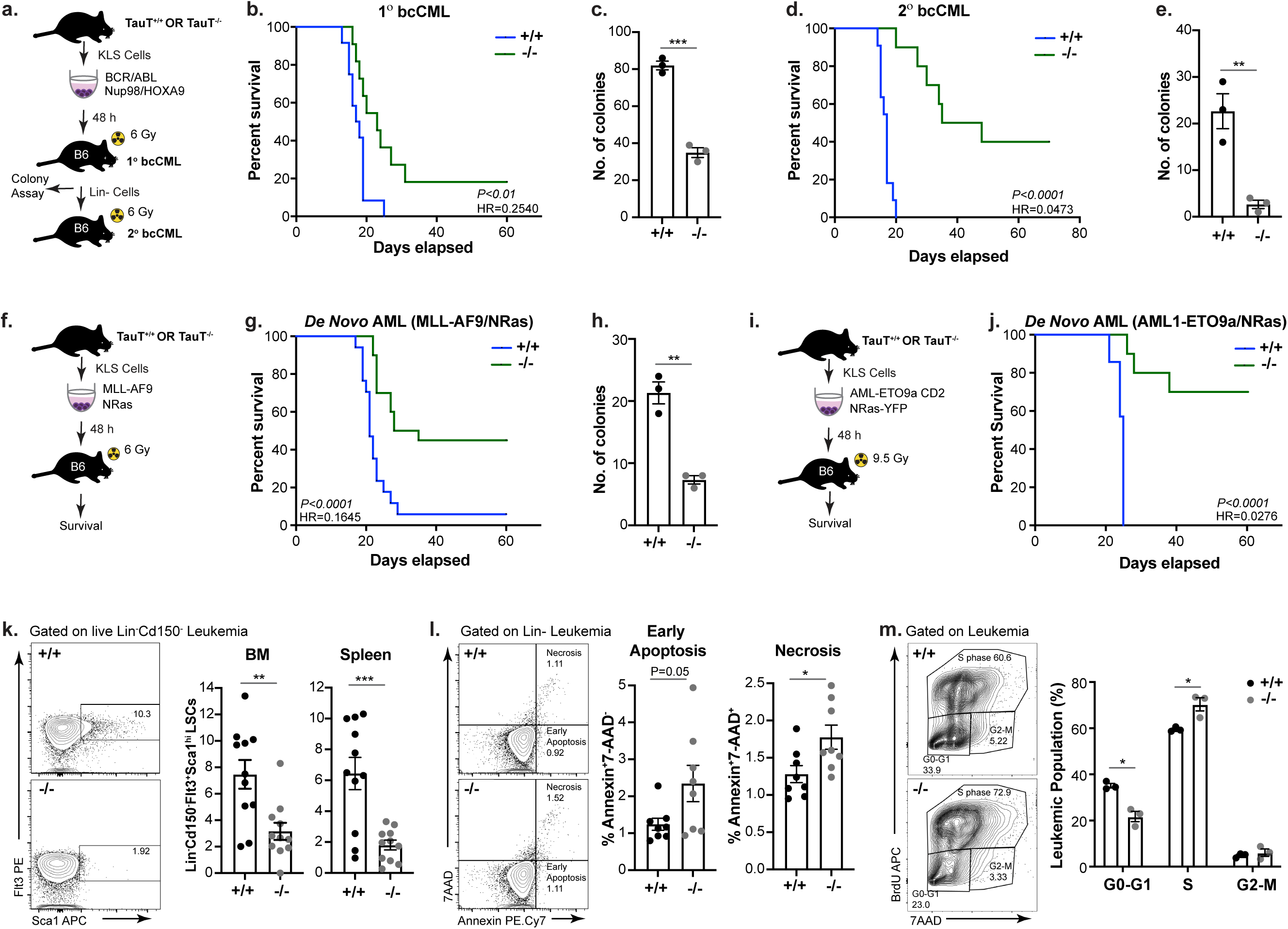
TauT loss impairs myeloid leukemia initiation and propagation in murine models. (a) Schematic shows the strategy used to determine the impact of TauT loss on bcCML initiation and LSC self-renewal. (b) Survival curve shows the impact of TauT loss on bcCML initiation *in vivo* in primary transplants. KLS cells from +/+ (TauT^+/+^) and -/- (TauT^-/-^) mice were retrovirally transduced with BCR-ABL and NUP98-HOXA9 and sorted transduced cells were transplanted (n=11-12 per cohort; data combined from three independent experiments; log-rank test). (c) Graph shows the colony-forming ability of Lin^-^ +/+ and -/- primary leukemia cells (n=3 replicates independent culture wells per cohort). (d) Lin^−^ leukemia cells were isolated from +/+ and -/- established cancers and transplanted in secondary recipients. Survival curve shows the impact of TauT loss on LSC self-renewal (n=10-11 per cohort; data combined from two independent experiments). (e). Colony forming ability of +/+ and -/- Lin^−^ cells from secondary transplants (n=3 replicates independent culture wells per cohort). (f) Schematic shows the strategy used to determine the impact of TauT loss on AML initiation. KLS cells from +/+ and -/- mice were retrovirally transduced with MLL-AF9 and NRas and transplanted in irradiated recipients. (g) Survival curve shows the impact of TauT loss on *de Novo* AML initiation (n=17 for +/+ and n=20 for -/-; data combined from four independent experiments; log-rank test). (h) Graph shows the colony-forming ability of cKit^+^ MLL-AF9^+^ NRas^+^ cells from +/+ and -/- established leukemia (n=3 replicates independent culture wells per cohort). (i) Experimental strategy to determine the impact of Taut loss on AML-ETO9a driven AML. (j) Survival curve of mice transplanted with KLS transduced will AML-ETO9a and NRas (n=7 for +/+ and n=10 for -/-; data combined from two independent experiments; log-rank test). (k) Representative FACS plots and graphs of Lin^−^CD150^−^Sca1^+^Flt3^+^ leukemia stem cells in +/+ and -/- bcCML. The frequency of these cells in the bone marrow (left) and the spleen (right) of recipient mice are shown (n=11 per cohort. Data combined from three independent experiments). (l) Representative FACS plots and graph show analysis of early and late apoptosis in bcCML cells 14 days post-transplant (n=8 per cohort; data combined from two independent experiments). (m) Representative FACS plots and graph show frequency of *in vivo* BrdU incorporation in +/+ and -/- bcCML. Average frequency of cells in distinct phases of the cell cycle are shown (n=3 per cohort). Error bars represent ±SEM; statistics from unpaired t-tests or as indicated. *p*<*0.05, **p*<*0.01, ***p*<*0.001

On a cellular level, TauT loss led to a 2.4 to 3.6-fold reduction in the primitive Lin^-^CD150^-^Flt3^+^Sca1^+^ bcCML stem cells (Fig. 4k)^7, 45^. Further, TauT loss increased apoptosis (Fig. 4l) and promoted cell proliferation (Fig. 4m). These data demonstrate a critical requirement for TauT in the initiation, self-renewal, and propagation of myeloid leukemia.

### TauT controls leukemia progression by modulating glycolysis

To determine downstream mechanisms controlled by TauT in myeloid leukemia progression, we analyzed wild-type and TauT^-/-^ leukemia cells by RNA-Seq (Fig. 5a) and identified 932 downregulated and 1158 upregulated genes in the absence of TauT (P_adj_<0.05). The 1158 upregulated genes (P_adj_<0.05) primarily constituted pathways associated with hematopoietic cell lineage and cell cycle regulation (Fig. S5a), consistent with our cellular level analysis showing reduced LSCs and increased cell proliferation in that absence of TauT (Fig. 4k-m). The 932 downregulated genes (P_adj_<0.05) primarily constituted pathways associated with glycolytic metabolism, including canonical glycolysis and glucose catabolic process to pyruvate (Fig. 5b-c, S5b-c). Given the strong correlation of TauT loss with impaired glycolysis, we determined the physiological changes in glycolysis rate and glycolytic activity using the Seahorse assay (Fig. 5d). Consistent with the striking loss in glycolysis associated genes in the absence of TauT (Fig. 5b-c), our experiments identified a 1.4-fold reduction in basal glycolysis and glycolytic capacity in TauT^-/-^ LSCs as compared to wild-type controls (Figs. 5d-e). Although we did not find any difference in basal respiration or respiratory capacity (Fig. 5f-g), the maximal oxygen consumption was 1.4-fold lower in TauT null cells in the presence of the mitochondrial uncoupler FCCP. It is probable that the striking defects in glycolysis may be inducing a concomitant decrease in maximal mitochondrial respiration. Collectively, our data indicate that glycolytic functions are likely the primary downstream effectors of TauT function.

**Figure 5:**
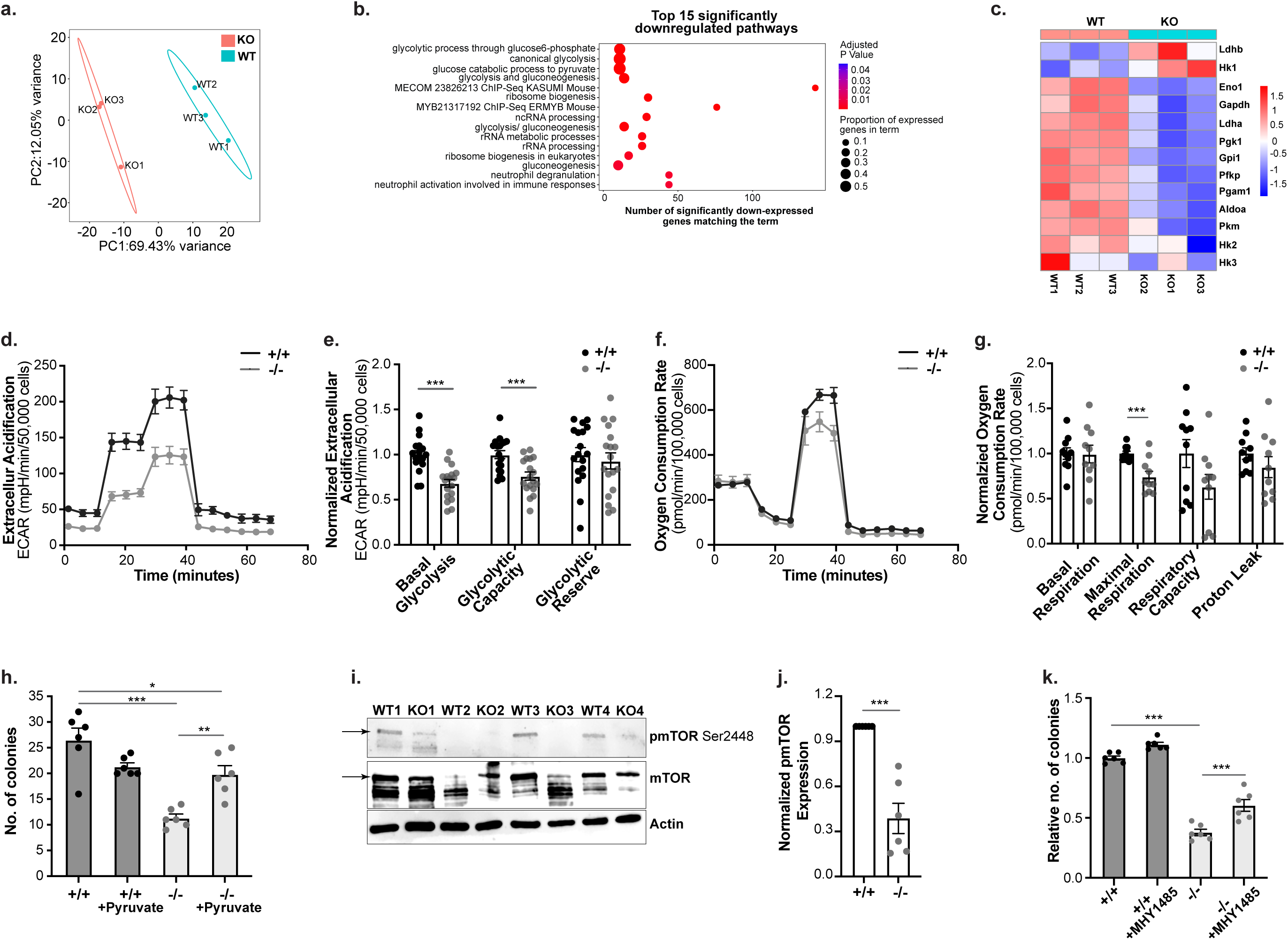
TauT loss impairs glycolytic metabolism in myeloid leukemia cells. (a) PCA-plot showing the distribution of the +/+ (TauT^+/+^) and -/- (TauT^-/-^) primary bcCML samples 11 days post-transplant (n=3 mice per cohort). (b) Enrichr analysis shows the top 15 pathways downregulated in -/- bcCML samples. (c) Heatmap shows expression of glycolysis associated genes in +/+ and -/- bcCML samples. (d) Representative curve of extracellular acidification (ECAR) in +/+ and -/- Lin-bcCML cells. (e) Quantification of changes in glycolysis, glycolytic capacity, and glycolytic reserve in -/- Lin-bcCML cells as compared to +/+ controls (n=4-5 independent culture wells per cohort; data combined from four independent experiments). (f) Representative curve of oxygen consumption rate (OCR) in +/+ and -/- Lin^-^ bcCML cells. (g) Normalized OCR in +/+ and -/- Lin^-^ bcCML cells (n=3-5 independent culture wells per cohort; data combined from three independent experiments). (h) Graph shows impact of supplementing pyruvate (1mM) on the colony forming ability of Lin^-^ bcCML cells (n=3 independent culture wells per cohort; data combined from two independent experiments; one-way ANOVA). (i) Immunoblot shows phospho-mTOR (pmTOR), mTOR, and Actin expression in 4 independent +/+ and -/- Lin^-^ bcCML samples (also see Supplementary Fig. S5d for additional blots). (j) Densitometric quantification of normalized pmTOR protein levels in +/+ and -/- Lin-bcCML cells (n=6 independent samples/cohort). (k) Colony-forming ability of +/+ and -/- Lin^-^ bcCML cells in the presence of the mTOR activator (MHY1485; 2μM) (n=3 independent culture wells per cohort; data combined from two independent experiments, one-way ANOVA). Error bars represent ±SEM; Statistics from unpaired t-tests or as indicated. *p<0.05, **p<0.01, ***p<0.001.

Since our data identify a profound reduction in glycolysis with TauT loss, we tested if bypassing glycolysis with pyruvate supplementation could rescue TauT defects. Our experiments showed that the colony forming ability of TauT^-/-^ LSCs could be significantly rescued in the presence of pyruvate, indicating that TauT dependent glycolytic functions are key drivers of disease progression (Fig. 5h). Given that mTOR signaling is known to play a key role in regulating expression of glycolysis related genes^46–48^, we tested if TauT loss impairs mTOR activation. Western blot analysis of LSCs showed a 2.6-fold loss in activated phosphorylated mTOR compared to controls (Figs. 5i-j, S5d). Importantly, the mTOR activator MHY1485^49^ could rescue the downstream expression of glycolysis associated genes (Fig. S5e) and the functional colony forming ability of TauT null LSCs to 60% of control (Fig. 5k). Collectively, our studies show that TauT regulates glycolysis in myeloid leukemia cells by activating mTOR signaling.

### TauT is essential for human AML growth and progression

Our RNA-Seq based expression analysis identified ∼2-fold higher *TAUT* expression in human bcCML and AML stem-enriched cells as compared to normal hematopoietic stem/progenitors (Fig. 2f), indicating that this gene may play a functional role in human disease progression. To test this, we determined the impact of inhibiting *TAUT* expression using shRNAs on the colony forming ability of human leukemia cell lines. Knocking down *TAUT* expression using two independent shRNAs (Fig. S6a) significantly impaired the colony forming ability of human bcCML (K562) and AML (THP1, MV-4-11) cell lines by 6 to 23.6-fold (Fig. 6a-c). Importantly, our experiments showed that three primary human AML patient samples carrying shRNAs against *TAUT* formed 2.33 to 9.12-fold fewer colonies *in vitro* (Fig. 6d). In contrast, we did not see any impact of *TAUT* inhibition on the colony forming ability of normal human CD34^+^ stem/progenitor cells from three independent donors (Fig. 6e). Finally, *TAUT* inhibition reduced the engraftment of primary human AML cells by 1.2 to 40-fold in patient derived xenograft models (Fig. 6f-g), indicating that *TAUT* expression and function is essential for human AML establishment *in vivo*.

**Figure 6:**
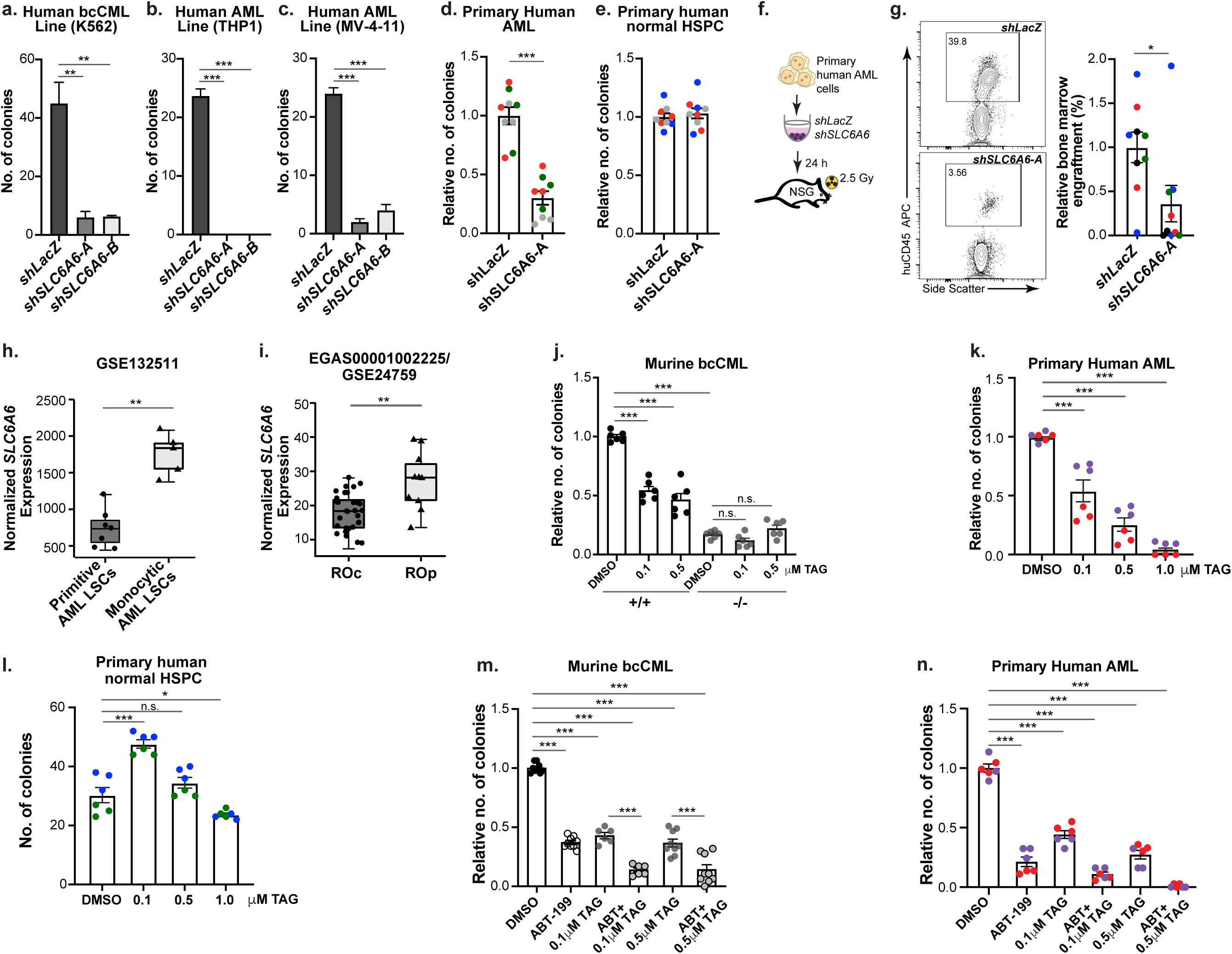
TauT inhibition impairs human leukemia growth and synergizes with venetoclax treatment. (a-c) Colony-forming ability of human leukemia cell lines transduced with lentiviral shRNAs targeting *LacZ* (control) or human *SLC6A6*, K562 (a), THP1 (b), and MV-4-11 (c; n=3 independent culture wells per cohort; one-way ANOVA). (d) Colony-forming ability of primary human patient-derived AML cells transduced with lentiviral shRNAs targeting *LacZ* (control) or human *SLC6A6* (each color represents an independent sample; n=3 independent culture wells per cohort; data combined from three independent experiments). (e) Colony-forming ability of 3 independent normal human CD34^+^ bone marrow hematopoietic stem cells and progenitors (HSPCs) transduced with lentiviral shRNAs targeting *LacZ* (control) or human *SLC6A6* (each color represents an independent sample; n=3 independent culture wells per cohort; data combined from 3 independent experiments). (f) Schematic shows the strategy used to determine the impact of TauT loss on human primary AML *in vivo*. (g) Representative FACS plots and graph show analysis of primary human AML bone marrow engraftment (CD45^+^) of xenograft recipients (Each dot represents a mouse, each color represents an independent sample; n=2-3 mice per sample). (h) Graph shows normalized *SLC6A6* expression in primitive and monocytic ROS-Low AML LSCs (median ± inter-quartile; n=7 for primitive and n=5 for monocytic; unpaired Wilcoxon’s test). (i) Graph shows normalized *SLC6A6* expression in relapse origin-committed (RO_C_) and relapse origin-primitive (RO_P_) AML (median ± inter-quartile; n = 29 for RO_C_, n=10 for RO_P_; unpaired Wilcoxon test). (j) Colonies formed by TauT^+/+^ (+/+) and TauT^-/-^ (-/-) bcCML Lin^-^ cells in the presence of DMSO (control) or indicated doses of taurine antagonist (TAG) (n=3 independent culture wells per cohort; data combined from two independent experiments; one-way ANOVA). (k) Colony forming ability of primary human AML cells in the presence of DMSO (control) or indicated doses of TAG (each color represents an independent sample; n=3 per sample; data combined from 2 independent experiments; one-way ANOVA). (l) Colony-forming ability of 2 independent normal human CD34^+^ bone marrow hematopoietic stem cells and progenitors (HSPCs) treated with indicated doses of TAG (each color represents an independent sample; n=3 per sample; data combined from two independent experiments; one-way ANOVA). (m) Graph shows relative colony forming ability of murine Lin^-^ bcCML cells in the presence DMSO (control), 500nM venetoclax, 0.1μM and 0.5 μM TAG or indicated combinations (n=3 per independent culture wells/treatment; data combined from 2-3 independent experiments; one-way ANOVA). (n) Relative colony forming ability of primary human AML cells in the presence of DMSO (control), 500 nM venetoclax, 0.1μM and 0.5 μM TAG or indicated combinations (each color represents an independent sample; n=3 per sample; data combined from two independent experiments; one-way ANOVA). Error bars represent ±SEM; statistics from unpaired t-tests or as indicated. *p*<*0.05, **p*<*0.01, ***p*<*0.001

Recent work indicates that monocytic AML subtypes can be resistant to the therapeutic Bcl-2 inhibitor, venetoclax^50^. Our analysis identified significantly higher *SLC6A6* expression in LSCs from monocytic AML as compared to primitive AML (Fig. 6h)^50^, and in relapsed AML originating from stem/progenitor like cells (RO_P_) as compared to more committed populations (RO_C_)^51^ (Fig. 6i), consistent with the strong correlation between SLC6A6 expression and poor prognosis (Fig. 2e). Given that venetoclax is known to impair oxidative phosphorylation in leukemia^9, 52^, we tested if genetic TauT loss can synergize with venetoclax to effectively impair LSC function. Our experiments showed a 3.2-fold loss in viability of TauT^-/-^ bcCML LSCs in the presence of venetoclax (ABT-199) as compared to controls (Fig. S6b). Importantly, while venetoclax alone reduced colony formation by 1.4-fold and TauT loss reduced this by 3.2-fold, their combination showed a striking 13.2-fold reduction in colonies as compared to wild-type controls (Fig. S6c).

To determine if a small molecule inhibitor of TauT can be effective in blocking leukemia cell growth, we used a previously described taurine antagonist (6-aminomethyl-3-methyl-4H-1,2,4-benzothiadiazine-1, 1-dioxide hydrochloride; TAG)^53, 54^. Mechanistically, the sulphonamide group in TAG mimics taurine to bind and block the activity of the taurine transporter^55^. Treating bcCML LSCs with TAG reduced intracellular taurine levels by 1.4-fold (Fig. S6d), indicating that it can block taurine uptake by leukemia cells. Further, TAG treatment led to a dose dependent reduction in the colony forming ability of wild-type murine bcCML Lin- and AML cKit^+^ leukemia stem-enriched cells but not TauT^-/-^ cells (Figs. 6j, S6e). Importantly, while TAG treatment impaired the growth of primary human AML cells in colony assays (Fig. 6k), it did not impair the colony forming ability of normal human CD34^+^ stem/ progenitors (Fig. 6l), similar to the effect of *TAUT* shRNAs (Figs. 6d-e).

Finally, we tested if TAG treatment can synergize with venetoclax in murine and human leukemia cells. Our experiments showed that while TAG or venetoclax alone impair colony formation by ∼2.5-fold, combining TAG with venetoclax reduces the colony forming ability of bcCML LSCs by 7.2-fold as compared to controls (Fig. 6m). To determine if TAG can also synergize with venetoclax to inhibit the growth of human leukemia cells, we tested the impact of TAG treatment in venetoclax resistant and sensitive AML cell lines^56^. While venetoclax treatment reduced the colony forming ability of the MV-4-11 cells by 2.2-fold, it had a minor impact on the resistant THP1 cells. However, combinatorial treatment with TAG and venetoclax led to a 1.6-fold reduction in the colonies formed by THP1 and 7.4-fold reduction in those formed by MV-4-11 (Figs. S6f-g) cells as compared to controls, indicating that TAUT inhibition can sensitize myeloid leukemia cells to venetoclax. Importantly, our experiments with primary patient-derived AML cells showed that while venetoclax reduces colony formation by 3.3 to 7.8-fold and TAG by 2 to 4.6-fold, combinatorial treatment profoundly impairs colony forming ability by 7.4 to 54.5-fold as compared to controls (Fig. 6n).

Collectively, our data identify a critical requirement for *TAUT* expression in primary human AML growth and provide a scientific rationale for combinatorial use TAUT inhibitors in the context of venetoclax.

## DISCUSSION

Our work here leverages single cell RNA-sequencing to define the changing landscape of the non-immune cancer microenvironment with disease progression, and to identify unique niche driven signals that promote disease progression. Single cell sequencing strategies have been effectively used to characterize the immune microenvironment of both solid tumors and leukemias^30, 57–62^. Although technical limitations of tumor dissociation processes and scRNA-seq based detection have hampered analyses of non-immune stromal cells, a few recent studies have identified antigen presenting cancer-associated fibroblasts and immune suppressive endothelial cells in lung and pancreatic cancers^63–65^. A reduction in osteoblasts has been noted during leukemia initiation in bone marrow niche^20^ damaged by extensive irradiation^48, 66^. However, dynamic alterations in the TME through the disease trajectory, especially in the context of aggressive myeloid leukemias, have not been defined.

Our temporal scRNA-Seq based analyses of the undamaged leukemia TME identifies an expansion in mesenchymal stromal cells and their immature osteo-associated progeny, along with a loss in mature osteo-associated cells. This skew in osteo-associated populations possibly results from the observed downregulation of MSC osteolineage differentiation signals during disease progression. Thus, our temporal gene-expression analysis of all osteo-associated populations in the TME may explain the conflicting findings using distinct candidate osteoblast markers in the leukemic niche^67, 68^. In addition, our data indicates that the temporal expansion in the arteriolar endothelial cells is accompanied with a loss in signals essential for endothelial cell integrity and function. These observations may explain the leaky blood vessels seen by *in vivo* imaging of AML bone marrow niche^1, 69^. In addition to population level changes, our studies also identify signals such as kit, thrombospondin, taurine and apolipoproteins from the TME that are essential for cancer progression. Importantly, while multiple TME ligands such as ApoE and Pvr can be detected all along the disease trajectory, the microenvironmental populations synthesizing them may change over time. Thus, therapeutic approaches aimed at targeting TME-driven signals may be more effective than blocking TME remodeling or inhibiting individual stromal populations.

Consistent with the clinicopathological similarities between AML and blast crisis CML^70, 71^, our gene expression analysis of primary human AML and bcCML CD34^+^ stem-enriched cells identifies distinct overlap in the two diseases. Our unbiased approach to determine LSC enriched cell surface receptors essential for disease progression discovered multiple genes known to be critical for myeloid cancers, including CD96^72^, CD47^73^, CD274^74^ and Protein Kinase D2^75^. Although all these genes are required for leukemic progression they are not associated with poor prognosis in AML (TCGA-LAML), unlike *LDLR* and *SLC6A6* that we describe here. It is thus likely that other cell surface receptors identified by our analysis also play a functional role in disease progression. Such candidates should be explored further to define new signals that may be of therapeutic significance in myeloid leukemias. Since the ApoE-LDLR and Taurine-TauT axes that we identify as critical for leukemic growth are known to play a role in aging^39, 76, 77^, it is possible that the cancer-associated signals in our TME-LSC interactome may be of broad relevance in aging-related disorders such as myelodysplastic syndromes.

Biosynthesis of taurine from cysteine is known to occur in liver, kidney, adipose tissues and pancreas^78^. Our data identify bone marrow osteolineage cells as a new source of taurine in the leukemia niche. While our *in vitro* studies blocking taurine production in the osteolineage cells establish a key role of TME-driven taurine synthesis in LSC survival and self-renewal, these data do not exclude the possibility that taurine, or β-alanine, produced outside the bone marrow niche can also contribute to disease progression. Since taurine is a common ingredient in energy drinks and is often provided as a supplement to mitigate the side-effects of chemotherapy^25^, our work defining a pro-cancer role for this amino acid indicates a need to re-evaluate the benefits of supplemental taurine in cancer patients.

Mechanistically, we identify a critical requirement of Slc6a6/TauT in regulating glycolysis in leukemia cells. While we note a minor effect of TauT loss on oxidative phosphorylation, the downstream effects of taurine transporter function in leukemia are primarily driven by changes in glycolytic capacity and can be rescued by circumventing glycolysis with pyruvate supplementation. Consistent with the observation that the widely used Bcl2-inhibitor, venetoclax, primarily targets oxidative phosphorylation in LSCs^9, 52^, we find significantly increased *SLC6A6* expression, and possibly glycolytic metabolism, in patient samples associated with venetoclax resistance. Our experiments showing a striking synergy between venetoclax and TauT inhibitors in venetoclax sensitive and resistant cell lines, as well as in primary human patient samples, indicate that combinatorial use of TauT inhibitors with venetoclax may be of therapeutic value in treating hematological cancers.

## AUTHOR CONTRIBUTIONS

B.J.R. and S.S performed majority of the experiments and helped write the paper; C.D.B. performed all bioinformatic analyses; C.M.K. provided experimental data and help; T.I. provided TauT knockout mice; J.L.L., L.M.C and M.W.B. provided primary leukemia patient samples and experimental advice; C.T.J. and J.M.A. carried out the human RNA-sequencing experiments and provided experimental advice; J.B. conceived of the project, planned and guided the research and wrote the paper.

## ACKNOWLEDGEMENTS

We are grateful to Erika Davidson and Jason Tran for technical support. We would like to thank Warren Pear and Ann Marie Pendergast for the BCR-ABL, D. Gary Gilliland for the NUP98-HOXA9, Christopher Counter for the NRasG12V, Scott Armstrong for the MLL-AF9, and Scott Lowe for the AML-ETO9a constructs. We would also like to thank the URMC Flow Cytometry Core for support with cell sorting. This work was supported by American Society of Hematology Scholar Award, Leukemia Research Foundation award and NIH grants R01DK133131 and R01CA266617 awarded to J.B.

## METHODS

### Generation of Experimental Mice

The *Slc6a6* (TauT) mice were bred as described earlier ^37^. For all murine leukemia experiments, the TauT^+/+^ and TauT2^-/-^ mice were used as donors and B6-CD45.1 (B6.SJL-*Ptprc^a^Pepc^b^/*BoyJ) or C57BL6/J mice were used as transplant recipients. For xenograft experiments with human cells, NSG mice (NOD.Cg-Prkdcscid Il2rgtm1Wjl/SzJ) were used as transplant recipients. All mice were 6–16 weeks of age. Mice were bred and maintained in the animal care facilities at the University of Rochester. All animal experiments were performed according to protocols approved by the University of Rochester’s Committee on Animal Resources.

### Cell Isolation and FACS Analysis

Cells were suspended in Hanks’ balanced salt solution (Gibco, Thermo Fisher Scientific) with 5% fetal bovine serum and 2 mM EDTA. Cells were prepared for FACS analysis and sorting as previously described ^1, 7^. Antibodies used for defining hematopoietic cell populations were as follows: CD3ε, CD4, CD8, Gr1, CD11b/Mac-1, TER119, CD45R/B220 and CD19 (all for lineage), CD117/cKit, Sca1, CD48 and CD150. All antibodies were purchased from, eBioscience (Thermo Fisher Scientific), BioLegend or BD Biosciences. Analysis was performed on LSRFortessa (Becton Dickinson), and cell sorting was performed on a FACSAria II (Becton Dickinson). Data was analyzed using FlowJo software.

### Retroviral and Lentiviral Constructs and Production

Retroviral MSCV-BCR-ABL-IRES-GFP (or -tNGFR) and MSCV-NUP98-HOXA9-IRES-YFP (or -huCD2 and -tNGFR) were used to generate bcCML. AML was generated with MSCV-MLL-AF9-IRES-tNGFR and MSCV-NRAS^G12V^-IRES-huCD2 or MSCV-AML-ETO9a and MSCV-NRAS^G12V^-IRES-YFP. Lentiviral short hairpin RNA (shRNA) constructs were designed and cloned in the pLV-hU6-EF1a-green or pLV-hU6-EF1a-red backbone (Biosettia) as per the manufacturer’s protocol. Virus was produced in 293T cells (ATCC) transfected with viral constructs along with VSV-G, Gag-Pol (retroviral production) or pRSV-rev, phCMV, and pMDlg/pRRE (lentivirus production) using X-tremeGENE-HP reagent (Roche). Viral supernatants were collected for three to six days followed by ultracentrifugal concentration at 20,000 rpm for 2 hr.

### Generation and Analysis of Leukemia Models

Bone marrow KLS cells were sorted from *TauT^+/^ or TauT^-/-^* mice and cultured overnight in X-VIVO15 (Lonza) media supplemented with 10% fetal bovine serum (Gemini), 50μM 2-mercatpoethanol, SCF (100ng/mL, R&D Systems), TPO (10ng/mL, R&D Systems) and Penicillin-Streptomycin (Gibco). Cells were retrovirally infected with MSCV-BCR-ABL-IRES-GFP (or -tNGFR) and MSCV-NUP98-HOXA9-IRES-YFP (or -huCD2 or - tNGFR) to generate blast crisis chronic myeloid leukemia. Acute myeloid leukemia was produced by sorting bone marrow KLS cells from *TauT^+/+^, or TauT^-/-^*mice and culturing in RPMI medium (Gibco, Thermo Fisher Scientific) supplemented with 20% fetal bovine serum, 50 μM 2-mercaptoethanol, 100 ng/mL SCF (R&D Systems), 10 ng/mL IL-3, and 10 ng/mL IL-6 (R&D Systems). Cells were retrovirally infected with MSCV-MLL-AF9-IRES-tNGFR and MSCV-NRAS^G12V^-IRES-YFP cells were harvested 48 hr after infection, sorted by FACs for BCR-ABL^+^, NUP98-HOXA9^+^ (bcCML only) and retro-orbitally transplanted into cohorts of sub-lethally irradiated (6 Gy) C57BL/6J mice. For AML-ETOa9 and NRAS, cells were transplanted into lethally irradiated (9.5 Gy) C57BL/6J recipients along with 3×10^5^ RBC lysed bone marrow rescue cells. For secondary transplants, Lin- cells from primary bcCML recipient mice were transplanted in secondary sub-lethally irradiated recipients. Recipients were maintained on acid water and sulfatrim diet and evaluated daily. Premorbid animals were euthanized, and relevant tissues harvested and analyzed by flow cytometry. Apoptosis assays were done using Annexin V and 7AAD (eBiosciences). Analysis of *in vivo* bromodeoxyuridine (BrdU) incorporation was performed using the APC BrdU Flow Kit (BD Biosciences) after a single intraperitoneal injection of BrdU (2 mg at 10 mg/mL).

### Isolation of Stromal Cells for RNA-Seq and Analysis

Stromal cells were isolated as described^79^. Briefly, bone and bone marrow (BM) were isolated from long bones and pelvis in 1x Media 199 (Gibco) with 2% fetal bovine serum (GeminiBio). BM was digested for 30 minutes in HBSS containing 2mg/mL Dispase II (Gibco), 1mg/mL Collagenase Type IV (Sigma-Aldrich), and 20ng/mL DNase Type II (Sigma-Aldrich). Bone spicules were digested for 60 minutes in PBS supplemented with 2.5mg/mL Collagenase Type I (Stem Cell Technologies) and 20% FBS. Digested bone marrow was RBC lysed using RBC Lysis Buffer (eBioscience). Bone and BM cells were pooled and CD45^+^ Ter119^+^ hematopoietic cells were magnetically depleted on an autoMACS cell separator (Miltenyi Biotec). The CD45-Ter119- stromal cells were either stained an analyzed for candidate populations by flow cytometry (BD LSRFortessa) or further enriched by sorting (BD FACSAria II) and processed for single cell RNA-Sequencing.

### scRNA-Sequencing and Analysis

Cell suspensions were processed to generate single-cell RNA-Seq libraries using Chromium Next GEM Single Cell 3′ GEM, Library and Gel Bead Kit v3.1 (10x Genomics), per the manufacturer’s recommendations. Samples were loaded on a Chromium Single-Cell Instrument (10x Genomics, Pleasanton, CA, USA) to generate single-cell GEMs (Gel Bead-in-Emulsions). GEM reverse transcription (GEM-RT) was performed to produce a barcoded, full-length cDNA from poly-adenylated mRNA. After incubation, GEMs were broken, the pooled GEM-RT reaction mixtures were recovered, and cDNA was purified with silane magnetic beads (DynaBeads MyOne Silane Beads, ThermoFisher Scientific). The purified cDNA was further amplified by PCR to generate sufficient material for library construction. Enzymatic fragmentation and size selection was used to optimize the cDNA amplicon size and indexed sequencing libraries were constructed by end repair, A-tailing, adaptor ligation, and PCR. Final libraries contain the P5 and P7 priming sites used in Illumina bridge amplification. Sequence data was generated using Illumina’s NovaSeq 6000. Samples were demultiplexed and counted using cellranger-4.0.0 mkfastq and count, using default parameters. The samples were aligned to a custom reference containing the 10X provided mm10-2020-A mouse reference and an additional EGFP sequence.

Seurat-4.1.0 within R-4.1.1 was used for most of processing. Additionally, dplyr-1.2.0 and tidyverse-1.3.2 were used extensively for data piping and transformation. Samples were imported and cells were filtered for at least 200 features captured per cell and features were filtered for expression in at least 3 cells. Additional filters were applied, filtering out all cells with higher then 5% mitochondrial content and filtering out all cells positive for CD45, CD71, Ter119, and EGFP. All samples were merged using ‘merge’ and normalized using SCTransform, regressing out the impact of mitochondrial features. Principal component analysis was performed using the first 40 principal components (RunPCA) and clusters were generated using FindNeighbors and FindClusters (resolution=0.5). UMAP dimensional reduction was also performed using RunUMAP using the first 30 principal components. Clusters were initially typed using scMCA-0.2.0. Populations typed as non- stroma (B-cell, Pre-B-cell, Pro-Erythrocyte, Pro-Erythroblast, Neutrophil, and Megakaryocyte) were filtered from the dataset. Populations were further typed using published bone marrow stroma markers^20^. FindAllMarkers was used within the context of specific populations to determine which genes change significantly over time. DEGreport-1.30.3 implemented degPatterns was used to tie specific expression patterns to the time points. Expression patterns corresponding to broadly increased or decreased expression over time were passed to gene set enrichment against KEGG_2019, GO_Biological_Process_2021, WikiPathways_2019_Mouse, and ChEA_2022 databases using EnrichR-3.0.

Ligand expression corresponding to bcCML and AML expressed receptors was determined via differential expression across the populations. Differentially expressed genes were filtered for corresponding ligands using a combination of nichenetr-1.1.0 included ligand-receptor mapping and literature supported interactions. These included Nrp1^80^, Adsl^81, 82^, Cdo1 and Csad for taurine synthesis^34^, Gadl1 and Cndp1 for β- alanine synthesis^35, 36^, Kit^83^, CD155^84^ and CD33^85^ for those not captured within this mapping. Broad Institute hosted dataset ^30^ was used as a stand in for healthy immune microenvironment within the context of AML. Counts (RNA_soupX1.mtx.gz) and metadata (metadata_clustering_w_header_upd.csv) were downloaded via the Broad Single Cell Portal and imported using Seurat. FindMarkers was used to determine which genes were up regulated within the microenvironment in relation to the other populations. Significantly expressed genes were then filtered for ligands using the same method as the stromal microenvironment. Significantly expressed ligand and receptor mappings within the stroma microenvironment, the immune microenvironment, and within bcCML and AML human bulkRNA were visualized using circlize-0.4.15. For illustration purposes, ligands expressed with a log fold change of 5 or higher were limited to 5.

### Primary Human Patient Derived Cells and Human Leukemia Cell Lines

For human RNA-Seq experiments and other studies, CD34^+^ cells were isolated from bone marrow samples of healthy donors and AML and blast crisis CML patient samples obtained under University of Rochester institutional review board-approved protocols with written informed consent in accordance with the Declaration of Helsinki. The normal human CD34^+^ HSPCs used in all functional assays were purchased (Stem Cell Technologies). Cells were cultured in Iscove’s modified Dulbecco’s medium with 10% FBS, 100 IU/mL Penicilin-Streptomycin (Gibco) and 55 μM 2-mercaptoethanol, 1x LDL (Sigma) and supplemented L-Gutamine with 100 ng/mL SCF and TPO (R&D Systems). Human leukemia cell lines K562, THP1, M-V-411 (ATCC) were maintained in RPMI with 10% FBS, 100 IU/mL Penicillin-Streptomycin (Gibco). For colony-forming assays with shRNAs, leukemia and normal cells were transduced with indicated lentiviral shRNAs. Cells were sorted 24 h post-infection and plated in CFU assays or transplanted in sub-lethally irradiated NSG mice.

### Primary Human CD34^+^ cells RNA-Sequencing Analysis

Total RNA was purified with Qiagen Rneasy PLUS kit per manufacturer recommendations and eluted in nuclease-free water. The total RNA concentration was determined with the NanopDrop 1000 spectrophotometer (NanoDrop) and RNA quality assessed with the Agilent Bioanalyzer (Agilent Technologies). The TruSeq Stranded mRNA Sample Preparation Kit (Illumina Inc.) was used for next generation sequencing library construction per manufacturer’s protocols. Briefly, mRNA was purified from 200ng total RNA with oligo-dT magnetic beads and fragmented. First-strand cDNA synthesis was performed with random hexamer priming followed by second-strand cDNA synthesis using dUTP incorporation for strand marking. End repair and 3’ adenylation was then performed on the double stranded cDNA. Illumina adaptors were ligated to both ends of the cDNA, purified by gel electrophoresis and amplified with PCR primers specific to the adaptor sequences to generate cDNA amplicons of approximately 200-500bp in size. The amplified libraries were hybridized to the Illumina single end flow cell and amplified using the cBot (Illumina Inc.). Single end reads of 100nt were generated for each sample using Illumina’s HiSeq2500v4.

Raw reads generated from the Illumina basecalls were demultiplexed using bcl2fastq v2.19.0. Quality filtering and adapter removal was performed using FastP v.0.20.1. Processed reads were then mapped to the human reference genome (hg38 + gencode version 36) using STAR_2.7.6a. Reads mapping to genes were counted using subread featurecounts v2.0.1. Differential expression analysis was performed using DESeq2-1.28.1 with a P-value threshold of 0.05 within R v4.0.2. Gene ontology analyses were performed using the EnrichR-3.0 package.

To determine significantly expressed receptors, the differential expression results were first filtered for significantly (padj<0.05) genes with a log fold change value of greater than 0. A list of potential cell surface receptors was generated using this gene list in conjunction with cell surface proteins detailed within the Cell Surface Protein Atlas ^28^ and essential for LSC growth *in vivo*^7^. The resultant gene list was further reviewed for genes not truly expressed within the cell surface, leading to the removal of CHST11, ST3GAL4, ACAA1, CLCN6, CHPF2, CTSK, ATP6AP1, CTSD, PNPLA6, DMXL2, TUBB6, MAN2B2, CLN3, MGAT4B, MYH9 and PIGG from further review.

### Taurine Quantification

Bone marrow cells or bone marrow peripheral fluid were isolated from femurs in Hanks’ balanced salt solution (Gibco) with 5% fetal bovine serum (GeminiBio) and 2mM EDTA. For taurine analysis in bone marrow cells, red blood cells were lysed with RBC lysis buffer (eBioscience) and suspended in RIPA buffer (Thermo Fisher Scientific) with benzonase nuclease (Sigma) and homogenized using a 0.5mL insulin syringe (BD Biosciences). The supernatant was concentrated using 10,000 MWCO spin columns (Corning). 25uL of concentrated samples were quantified using Taurine Assay kit (Sigma-Aldrich) and absorbance was measured at 415nm.

### MSC Isolation, Osteogenic Differentiation, and Co-culture with Leukemia Cells

Mesenchymal stromal cells (MSCs) were isolated from leukemic mice and cultured in 10cm dishes in MEM α with no ascorbic acid (Gibco) supplemented with 15% FBS and 100 IU/mL Penicillin/Streptomycin (Gibco). 6 days post culture initiation, the cells were sorted for CD45^-^CD3^-^B220^-^Ter119^-^Gr-1^-^CD31^-^Cd51^+^Sca1^+^ MSCs. Sorted cells were expanded in media descried above. For co-culture experiments, 50,000 MSCs were plated in a 48 well plate, and transduced with indicated lentiviral shRNAs. Three days post-infection, osteogenic differentiation was induced by switching to MEM α (Gibco) supplemented with 50μg/ml ascorbic acid (Sigma-Aldrich), 10mM β-glycerolphosphate (Sigma-Aldrich) and 100nM Dexamethasone (Sigma-Aldrich) for 6 days. Day 6 post-differentiation, 100,000 Lin- bcCML cells were added in X-Vivo supplemented with 10% FBS, 50μM 2-mercatpoethanol and Penicillin-Streptomycin. 3 days post co-culture, leukemia cells were analyzed for cell viability and plated in methylcellulose for colony forming assays.

### Normal HSC In Vivo Transplantation Assays

For bone marrow transplants, 500 HSCs were isolated from bone marrow of TauT^+/+^ or TauT^-/-^ mice and transplanted into lethally irradiated (9.5 Gy) CD45.1 mice along with 2×10^5^ Sca1-depleted bone marrow rescue cells. For subsequent secondary transplants, 2x10^6^ RBC lysed bone marrow cells isolated from primary recipient mice were transplanted into lethally irradiated (9.5 Gy) CD45.1 mice. Peripheral blood of recipient mice was collected every 4 weeks for 4 months post transplant and bone marrow analyzed at the end of 4 months.

### Methylcellulose Colony Formation Assays

For colony assays with mouse cells, indicated numbers of Lin- bcCML cells or cKit+ cells were plated in methylcellulose medium (M3234, StemCell Technologies). Colonies were scored at 7d. Colony assays with human cell lines or patient derived AML samples, cells were plated in complete methylcellulose medium (H4434 StemCell Technologies). Colony numbers were counted 10–14 days after plating. Taurine antagonist or TAG (synthesized by Enamine Inc.; 92% pure), mTOR activator (MHY1485; Sigma-Aldrich) and Venetoclax (ABT-199; Tocris Bioscience) were used as described.

### Seahorse Assay

Seahorse XF Glycolysis Stress Test Kit (Agilent Technologies, 103020-100) and Seahorse XF cell mito stress test kit (Agilent Technologies, 103015-100) were used to measure glycolytic flux (ECAR) and oxygen consumption (OCR) respectively. Murine Lin^-^ bcCML cells were sorted and cultured for 48 hours in X-Vivo supplemented with 10% FBS, SCF and TPO. One hour prior to analysis, 50,000-100,000 cells were seeded in a Cell-Tak (Corning, 324240) coated 96-well XF96 well plates in seahorse XF media (Agilent Technologies, 102353-100), and the plate was incubated at 37°C. ECAR data was measured following sequential addition of glucose (10mM), oligomycin (1μM) and 2-deoxyglucose (50mM) using XF96 analyzer (Agilent Technologies). OCR data was measured following sequential addition of oligomycin (1.5μM), Carbonyl cyanide-4 (trifluoromethoxy) phenylhydrazone (FCCP) (0.5μM), Rotenone and Antimycin (0.5μM) using XF96 analyzer.

### Western Blot Analysis

Cell lysates prepared in 1x RIPA (Thermo Scientific) supplemented with 1x protease, 1x phosphatase inhibitors (Cell Signaling Technology) and 250IU Benzonase Nuclease (Millipore Sigma) were separated on gradient polyacrylamide gels and transferred to nitrocellulose blotting membrane (0.45uM; GE healthcare Primary antibodies against phospho mTOR (Cell signaling), total mTOR (Cell signaling) and β-actin (Cell signaling) were used. Horse-radish peroxidase (HRP)-conjugated anti- rabbit antibody (Cell signaling) were used to detect primary antibodies. Immunoblots were developed using SuperSignal West Femto Maximum Sensitivity Substrate (Thermo Scientific).

### Murine TauT WT and KO Leukemia RNA-Sequencing

The Rneasy Plus Micro Kit (Qiagen, Valencia, CA) was used for RNA extraction. RNA concentration was determined with the NanopDrop 1000 spectrophotometer (NanoDrop) and RNA quality assessed with the Agilent Bioanalyzer 2100 (Agilent Technologies). 1ng of total RNA was pre-amplified with the SMARTer Ultra Low Input kit v4 (Clontech) per manufacturer’s recommendations. The quantity and quality of the subsequent cDNA was determined using the Qubit Flourometer (Life Technnologies) and the Agilent Bioanalyzer 2100 (Agilent). 150pg of cDNA was used to generate Illumina compatible sequencing libraries with the NexteraXT library preparation kit (Illumina Inc.) per manufacturer’s protocols. The amplified libraries were hybridized to the Illumina flow cell and sequenced using the llumina NextSeq 550 (Illumina Inc.). Single end reads of 100nt were generated for each sample. The mouse bulk RNA-Seq samples were processed otherwise identically to the human bulk RNA-Seq with two exceptions: m38 + gencode M27 reference for use within alignment and counting, and “-s 0” being used within subread featureCounts.

### Statistical Analysis

Statistical analyses were carried out using Graphpad Prism software version 6.0 (GraphPad software Inc.). Data are mean ± SEM. One-way ANOVA, Student’s t-tests, and Log-Rank tests were used to determine statistical significance.

## Supplementary Figure Legends

**Figure S1: Temporal changed in the leukemia bone marrow microenvironmental populations.**

(a) Survival curve shows trajectory of bcCML progression in unirradiated recipients. The time course for the scRNA-seq experiment (naïve, initiation, expansion, and end) is indicated (n=13, data combined from three independent experiments). (b) Line graphs show temporal changes in the proportion of all major lineages, and in the sub-clusters of chondrocyte and fibroblast populations. (c) Representative FACS plots and graph show changes in bone marrow sinusoidal and arteriolar endothelial cell frequency over time (n=3 mice per timepoint). (d) Gene clusters associated with changes in MSC, chondrocyte, and arteriolar endothelial populations during disease progression. (e) Enrichr plots show top ten upregulated and downregulated pathways in sinusoidal and arteriolar endothelial cells, as well as in MSCs and osteo-lineage populations.

**Figure S2: Temporal Changes in Leukemia Bone Marrow Microenvironmental Signals**

(a) Overlap between genes upregulated in bcCML and AML CD34^+^ cells compared to normal CD34+ cells, proteins expressed on cell surface^28^ and those that drop-out by 2-fold or more in the leukemia *in vivo* CRISPR screen^7^ (n=7 BM, n=10 bcCML, n=11 AML). (b) Average expression of ligands for cell surface proteins identified in Fig. 2b in primary human adult and pediatric AML cells, human AML immune microenvironment populations, and in normal cells^30^. (c-f) UMAP plot of gene expression within populations over time (naïve=0 days, initiation=2-4 days, expansion=7-9 days and, end=11-14 days post-transplant) with indicated changes in gene expression patterns.

**Figure S3: Impact of Inhibiting Leukemia Bone Marrow Microenvironmental Signals**

(a) Normalized *LDLR* expression in CD34^+^ cells from human bcCML, AML or normal BM (n=7 BM, n=10 bcCML, n=11 AML; Significance determined by deSeq2). (b) Experimental strategy used to determine the impact of MSC ApoE inhibition on co-cultured bcCML Lin- cells. (c) Relative *Apoe* expression in MSCs transduced with shRNAs targeting *LacZ* (control) or *Apoe* (n=3 technical replicates per cohort). (d) Number of live leukemia cells 72h post co-culture with MSCs transduced with *shApoe* or *shLacZ* (e) Colony forming ability of leukemic cells co-cultured for 72h with MSCs transduced with *shApoe* or *shLacZ* (n=3-4 independent culture wells per cohort; Data combined from two independent experiments). (f) Taurine biosynthesis pathway. (g) UMAP plot of *Csad* during disease progression (naïve=0 days, initiation=2-4 days, expansion=7-9 days and, end=11-14 days post-transplant). (h) Pictomicrographs of alizarin red staining of calcium deposits in primary murine bone marrow MSC before induction of osteogenic differentiation (Day 0) and at indicated times. (i) Relative expression of *Cdo1* and *Csad* in MSCs undergoing osteogenic differentiation (n=3 technical replicates per cohort; one-way ANOVA). (j) Taurine levels in MSC culture media during osteogenic differentiation, corrected for taurine amounts in unconditioned fresh media (n= 3 technical repliates per cohort; data shows taurine secreted over 48-72h as media is changed every 2-3 days; one-way ANOVA). (k) Relative *Cdo1* expression in MSCs transduced with shRNAs targeting *LacZ* (control) or *Cdo1* (n=3 technical replicates per cohort). Error bars represent ±SEM; Statistics from unpaired t-tests or as indicated. *p*<*0.05, **p*<*0.01, ***p*<*0.001

**Figure S4: The role of TauT in normal hematopoietic stem cell function.**

The frequency of (a) HSCs (KLSCD150^+^CD48^-^) and multipotent progenitors (MPPs KLSCD150^-^CD48^-^), (b) committed progenitors (granulocyte–macrophage progenitor (GMP), Lin^−^IL7Ra^−^Kit^+^Sca1^−^CD34^+^CD16/3^+^; common myeloid progenitor (CMP), Lin^−^IL7Ra^−^Kit^+^Sca1^−^CD34^+^CD16/32^−^; megakaryocyte–erythroid progenitor (MEP) Lin^−^IL7Ra^−^Kit^+^Sca1^−^CD34^−^CD16/32^−^), (c-d) differentiated hematopoietic cells in the bone marrow of TauT^+/+^ (+/+) and TauT^-/-^ (-/-) mice (n=5 mice per cohort; data combined from four independent experiments). (e-j) Total count of white blood cells and lymphocytes (e), granulocytes (f), monocytes (g), platelets (h), red blood cells (i), and hemoglobin (j) content in age and sex-matched 8 week old littermates (n=11 for +/+ and n=13 for -/-). (k) Average donor chimerism in the bone marrow of primary recipients, four months post HSC transplant (n=9 for +/+ and n=10 for -/-; data combined from two independent experiments). (l-n) Frequency of KLS (Lin^-^cKit^+^Sca^+^), HSCs, and MPPs (l), committed progenitors GMP, CMP, and MEP (m) and, differentiated hematopoietic (n) cells in bone marrow of primary HSC transplant recipients, four months post-transplant (n=9 for +/+ and n=10 for -/-; data combined from two independent experiments). (o) Average donor chimerism in the bone marrow of secondary bone marrow transplant recipients, four months post-transplant (n=6 per cohort). (p-s) Total frequency of KLS, HSCs, and MPPs (p), committed progenitors GMP, CMP, and MEP (q) and differentiated hematopoietic cells (r-s) in the bone marrow of secondary transplant recipients (n=6 per cohort). Error bars represent ±SEM; Statistics from unpaired t-tests. *p*<*0.05, **p*<*0.01, ***p*<*0.001

**Figure S5: Impact of TauT loss on energy metabolism pathways in myeloid leukemia.**

(a) Top 20 upregulated pathways in TauT^-/-^ (-/-) time matched bcCML as compared to TauT^+/+^ (+/+) controls. (b) Schematic shows genes regulating glycolysis. (c) Relative expression of glycolysis associated genes in +/+ and -/- Lin^-^ bcCML cells (n=3 technical replicates per cohort). (d) Western blot shows phospho-mTOR (pmTOR), mTOR, and Actin protein expression in 3 independent +/+ and -/- Lin^-^ bcCML samples. Also see Fig. 5g for additional blots. (e) Relative expression of glycolysis related genes in Lin^-^ bcCML cells from +/+ and -/- mice treated with 0.2μM MHY1485 for 48 h (n=3 technical replicates per cohort; data combined from 2 independent experiments). Error bars represent ±SEM; Statistics from unpaired t-tests. *p*<*0.05, **p*<*0.01, ***p*<*0.001

**Figure S6: A Small Molecule Taurine Antagonist Synergized with Venetoclax Treatment**

(a) Relative *SLC6A6* expression in K562 cells transduced with shRNAs targeting *LacZ* (control) and *SLC6A6* (n=3 technical replicates per cohort). (b) Viability of murine +/+ (TauT^+/+^) and -/- (TauT^-/-^) Lin^-^ bcCML cells treated with 500nM venetoclax (ABT-199) for 48 hours (n=3 independent culture wells per cohort; Data combined from 2 independent experiments; one-way ANOVA) (c) Impact of venetoclax (500nM) treatment on the colony forming ability of murine +/+ and -/- Lin^-^ bcCML cells (n=3 independent culture wells per cohort; data combined from 3 independent experiments; one-way ANOVA). (d) Graph shows taurine levels in lysate of +/+ bcCML leukemia stem cells treated with DMSO (control) or 0.5μM TAG (n=6; data combined from 2 independent experiments). (e) Colonies formed by ckit^+^ AML stem cells in the presence of DMSO (control) or indicated doses of TAG (n=3 independent culture wells per cohort; one-way ANOVA). (f) Colony forming ability of venetoclax sensitive MV-4-11 cells in the presence of venetoclax (ABT-199; 100nM) and indicated doses of TAG (n=3 independent culture wells per cohort; one-way ANOVA). (g) Colony forming ability of venetoclax resistant THP1 cells in the presence of venetoclax (ABT-199; 100nM) and indicated doses of TAG (n=3 independent culture wells per cohort; one-way ANOVA). Error bars represent ±SEM; statistics from unpaired t-tests or as indicated. *p*<*0.05, **p*<*0.01, ***p*<*0.001

