## Supplementary Figure 1 for "Temporal Single Cell Analysis of Leukemia Microenvironment Identifies Taurine-Taurine Transporter Axis as a Key Regulator of Myeloid Leukemia"

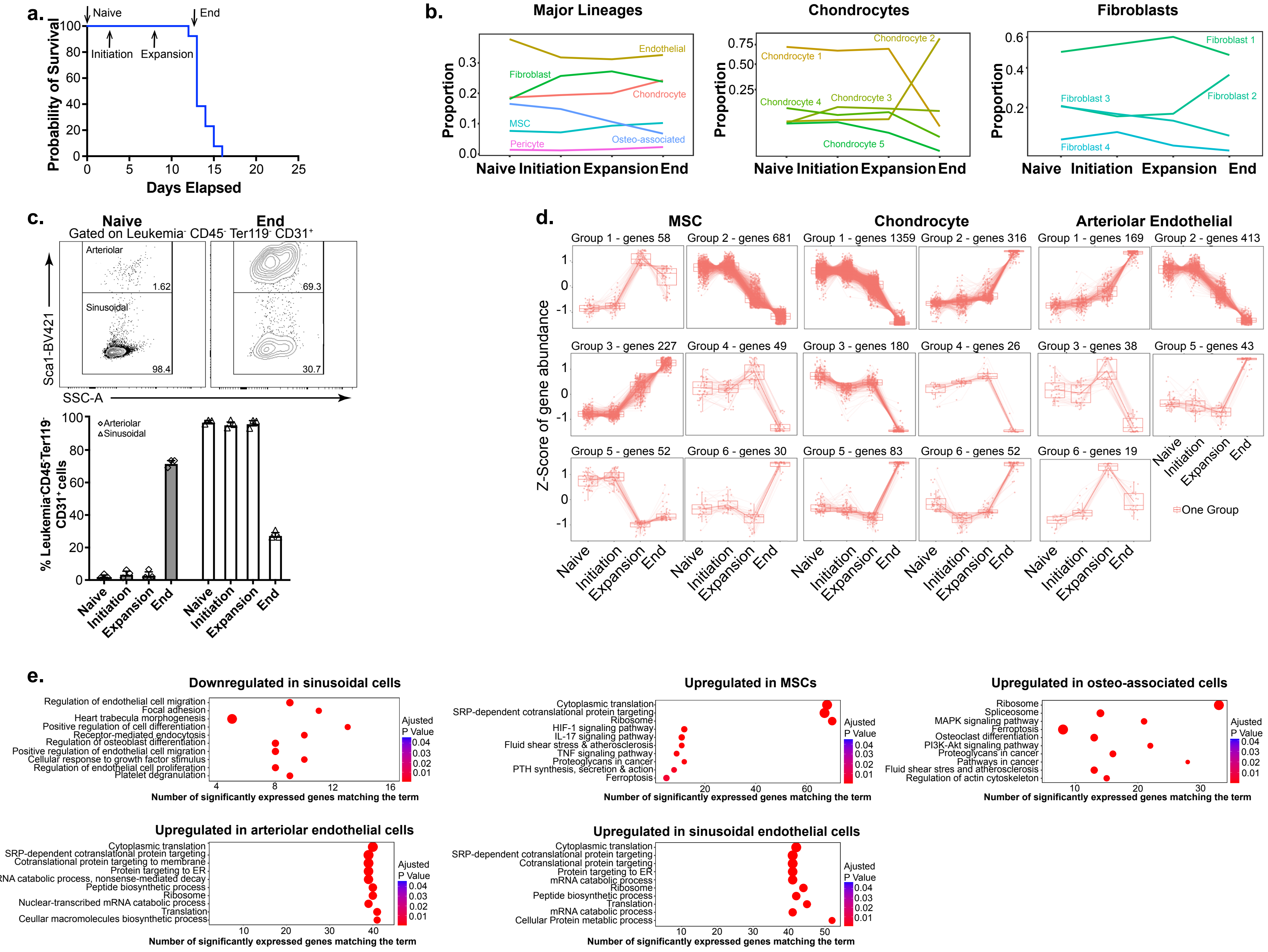


**Supplementary Figure S1: Temporal Changes in the Leukemia Bone Marrow Microenvironmental Populations**
