## Supplementary Figure 2 for "Temporal Single Cell Analysis of Leukemia Microenvironment Identifies Taurine-Taurine Transporter Axis as a Key Regulator of Myeloid Leukemia"

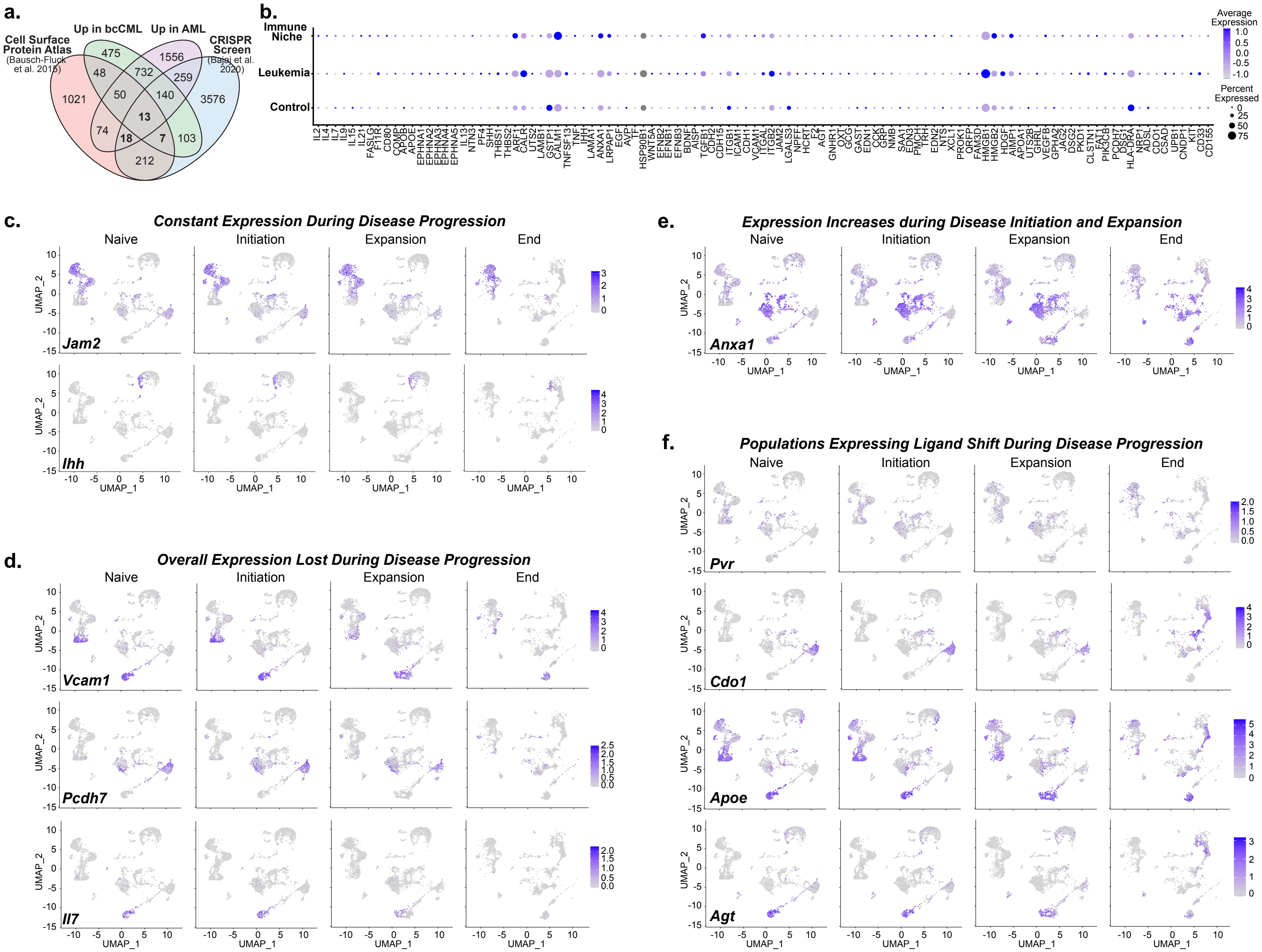


**Supplementary Figure S2: Bone Marrow Microenvironmental Interactions with Leukemia Cell Surface Receptors**
