## Supplementary Figure 3 for "Temporal Single Cell Analysis of Leukemia Microenvironment Identifies Taurine-Taurine Transporter Axis as a Key Regulator of Myeloid Leukemia"

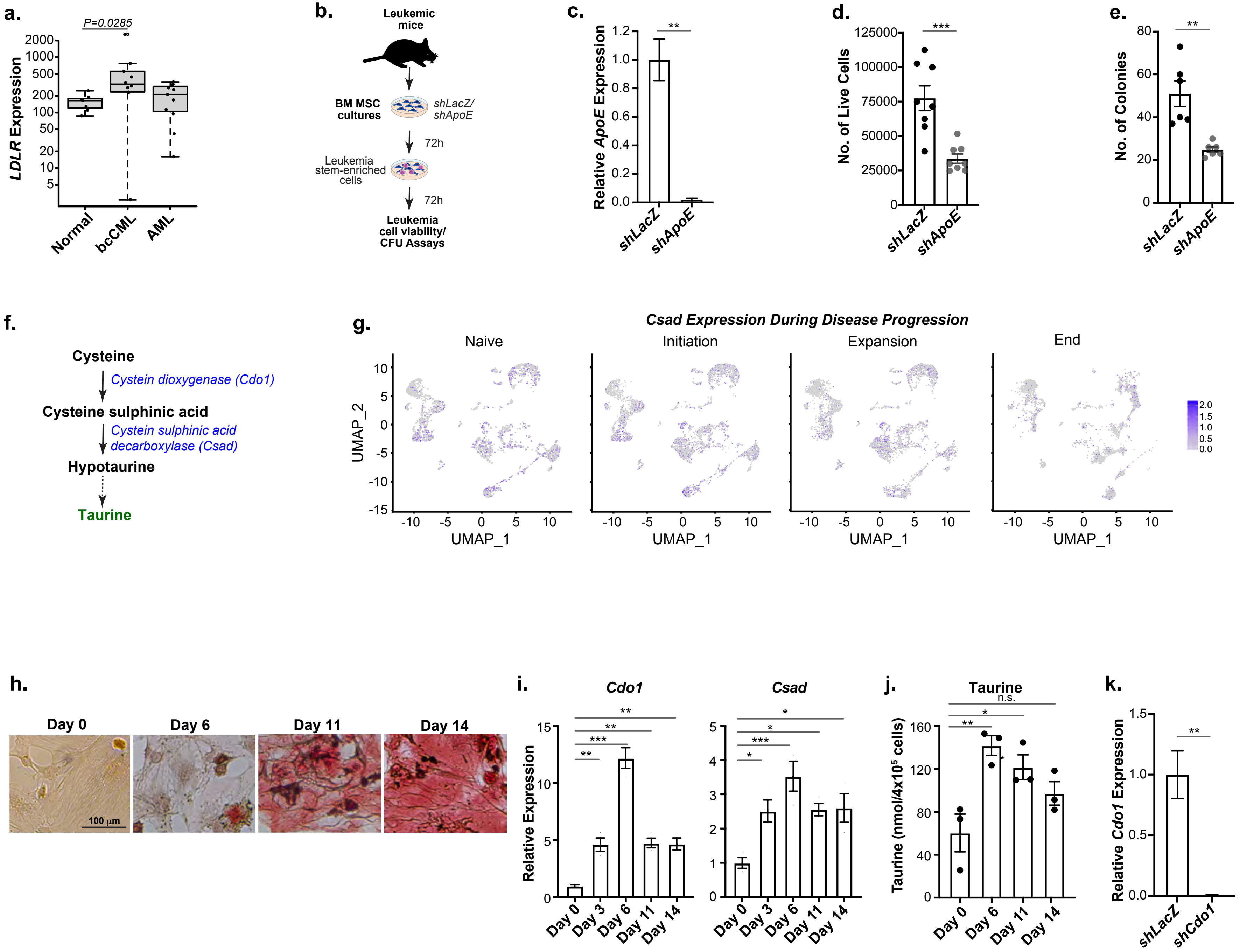


**Supplementary Figure S3: Impact of Inhibiting Leukemia Bone Marrow Microenvironmental Signals**
