## Supplementary figures and images for "Temporal Single Cell Analysis of Leukemia Microenvironment Identifies Taurine-Taurine Transporter Axis as a Key Regulator of Myeloid Leukemia"

### Supplementary Figure 4

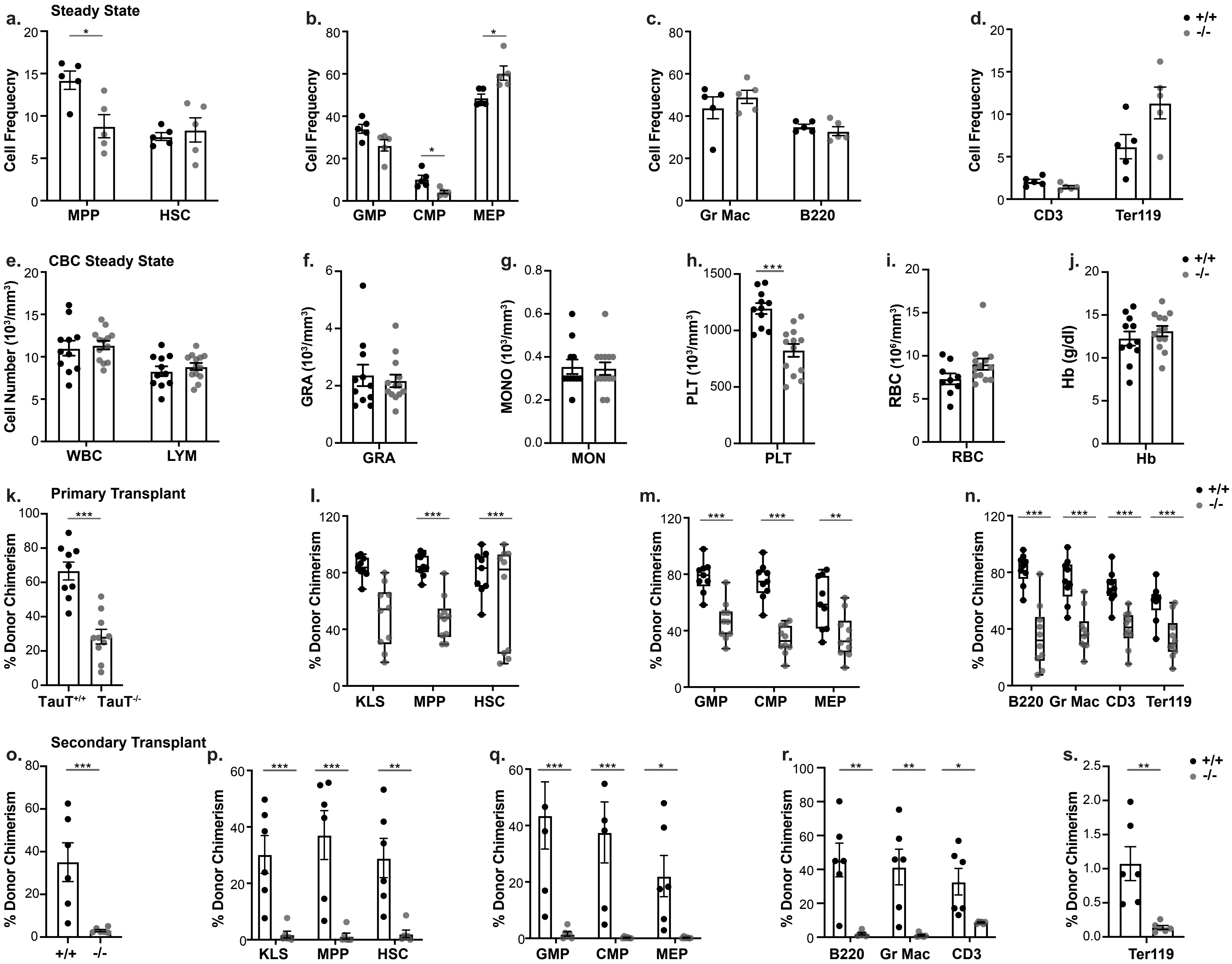


**Supplementary Figure S4: The Role of TauT in Normal Hematopoietic Stem Cell Function**
