## Supplementary Figure 5 for "Temporal Single Cell Analysis of Leukemia Microenvironment Identifies Taurine-Taurine Transporter Axis as a Key Regulator of Myeloid Leukemia"

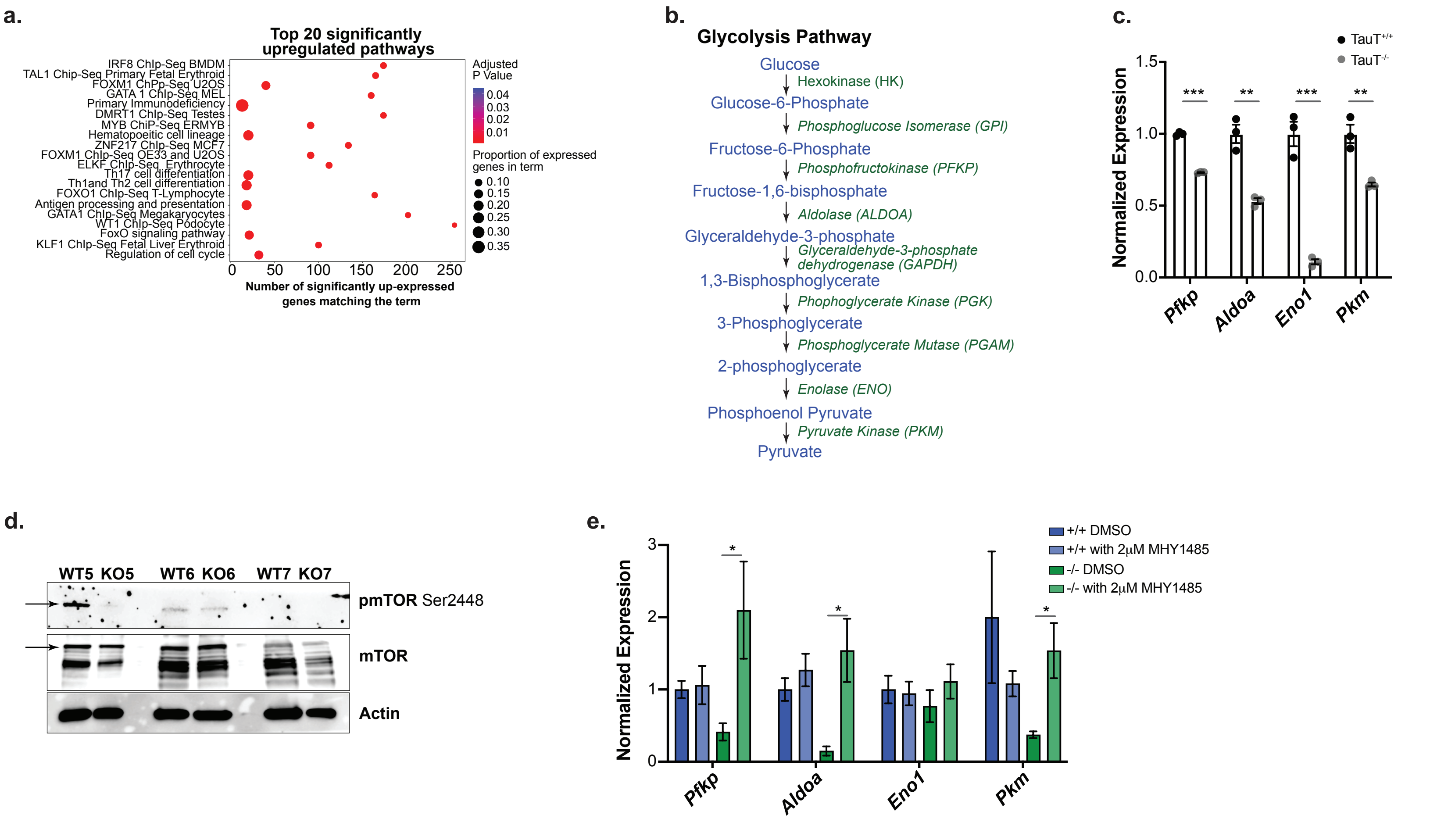


**Supplementary Figure S5: Impact of TauT Loss on Energy Metabolism Pathways in Myeloid Leukemias**
