## Supplementary Figure 6 for "Temporal Single Cell Analysis of Leukemia Microenvironment Identifies Taurine-Taurine Transporter Axis as a Key Regulator of Myeloid Leukemia"

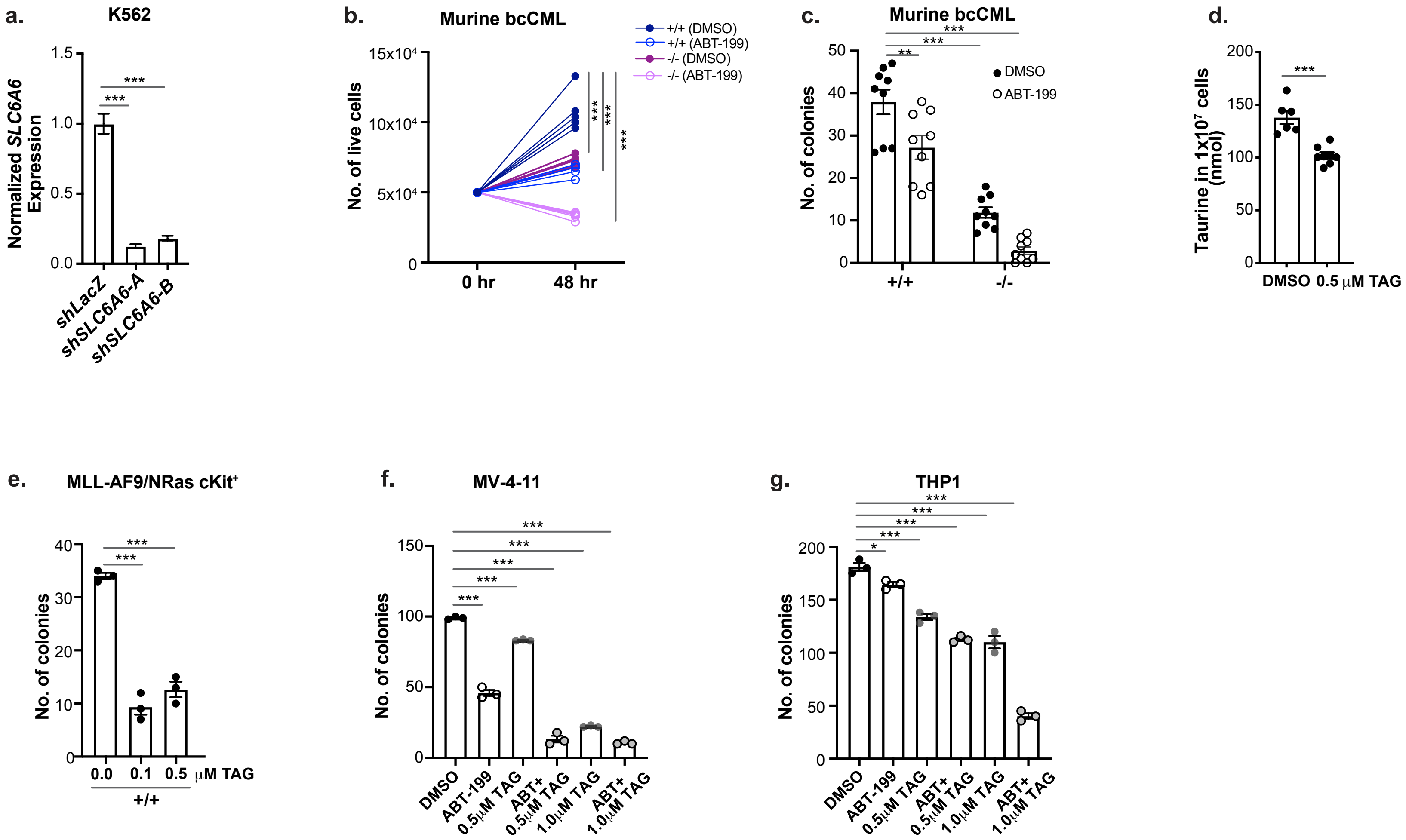


**Supplementary Figure S6: A Small Molecule Taurine Angtagonist Synergizes with Venetoclax Treatment**
